# SPECTRA: Spatial Inference for Tractometry Toward Precision Mapping of White Matter Microstructure

**DOI:** 10.64898/2026.05.08.723622

**Authors:** Yixue Feng, Julio E. Villalón-Reina, Iyad Ba Gari, Jonathan Davis Alibrando, Sophia I. Thomopoulos, Kenny Liou, Sunanda Somu, Hannah Yoo, Yuhan Shuai, Sasha Chehrzadeh, Talia M. Nir, Neda Jahanshad, Bramsh Qamar Chandio, Paul M. Thompson

## Abstract

Diffusion MRI tractometry characterizes white matter microstructure along fiber bundles, but standard along-tract profiling collapses measurements across the bundle cross-section, obscuring radial heterogeneity and producing spatially inconsistent units of inference. We present SPECTRA (Spatial Inference for Tractometry), a framework designed to address these limitations through a unified design of parameterization and statistical inference. First, we propose a 2D bundle parameterization that extends along-tract profiling to include a radial dimension defined on the atlas bundle. Second, we develop a two-stage hierarchical false discovery rate (hFDR) procedure for multi-bundle inference, which aggregates evidence at a coarser spatial scale before proceeding to finer-grained inference, with spatial scales derived from a Matérn kernel. Across extensive simulation conditions, we found that hFDR improves statistical power and reduces the sample size required to detect effects compared to global FDR correction, while maintaining appropriate error control. We further characterized how sensitivity-specificity tradeoffs depend on sample size, the magnitude, spatial extent, and configurations of effects, thereby providing practical guidance for tractometry study design. In an empirical analysis of mild cognitive impairment and dementia in more than 4,000 subjects across 63 bundles, SPECTRA revealed spatially localized patterns that were absent in 1D profiles. Together, these results demonstrate that spatially resolved parameterization and adaptive error control jointly enable precise mapping of white matter microstructure in large-scale tractometry studies. SPECTRA is openly available as a Python package.

## 1 Introduction

Diffusion MRI (dMRI) enables non-invasive quantitative evaluation of the brain’s white matter (WM) microstructure by measuring the directional diffusivity of water in neural tissue (Le Bihan et al., 1986). Derived microstructural measures, such as fractional anisotropy (FA) and mean diffusivity (MD) from diffusion tensor imaging (DTI) (Le Bihan et al., 2001), serve as indirect markers of local tissue properties, including axonal integrity and myelination. Many group-level analyses of WM microstructure pool measures across broad regions of interest (ROI) given a reference atlas (Jahanshad et al., 2013). Voxel-based analysis of dMRI offers even finer spatial resolution, but produces a large number of tests and is vulnerable to local misalignment of the WM structure across subjects (J. E. Lee et al., 2009). Tract-based Spatial Statistics (TBSS) (Smith et al., 2006) addresses misalignment by projecting subject data onto a common WM skeleton derived from FA.

Recent advances in tractography have enabled subject-specific segmentation of WM bundles that can preserve individual variability and establish cross-subject correspondence via a reference atlas (Garyfallidis et al., 2018; Schilling et al., 2025). Tractometry (Bells et al., 2011; Jones et al., 2005) then characterizes microstructural measures along the trajectory of each bundle at a spatial resolution finer than ROIs but coarser than individual voxels. Such along-tract mapping has revealed spatially localized profiles of microstructural abnormalities in bipolar disorder (Nabulsi et al., 2023), autism spectrum disorder (Kim et al., 2025), and Alzheimer’s disease (B. Q. Chandio et al., 2022; Dou et al., 2020; Feng et al., 2024), and has helped to identify links between amyloid and tau pathology and along-tract abnormalities (B. Q. Chandio et al., 2024b, 2025). Existing tractometry frameworks, including Automated Fiber Quantification (AFQ)(Yeatman et al., 2012), Tracts Constrained by UnderLying Anatomy (TRACULA) (Yendiki et al., 2011), Bundle Analytics (BUAN) (B. Q. Chandio et al., 2020), and Medial Tractography Analysis (MeTA) (Ba Gari et al., 2023, 2025) adopt a one-dimensional (1D) along-tract parameterization where each bundle is divided into a fixed number of segments/nodes along its principal axis.

While 1D profiling is well-suited for compact and elongated bundles such as the fornix or uncinate fasciculus, they are not sufficient to represent geometrically more complex structures such as the corpus callosum or thalamic radiations, which have large cross-sections. Several works have proposed a two-dimensional (2D) parameterization of WM bundles by fitting continuous medial representations (Yushkevich et al., 2008) and principal surfaces (Yue et al., 2016). Both methods anchor the parameterization in a fitted geometric surface, and typically require data of consistent quality across subjects, so that a surface can be reliably fitted. To ensure robust and efficient parameterization in data of variable quality while preserving cross-subject biological variability in the reconstructed bundles, we extend the streamline-based 1D parameterization of BUAN and define a 2D coordinate system directly on the atlas bundle streamlines, with user-specified resolutions along- and radially across the bundle. This produces a common grid that enables cross-subject correspondence and local measures of microstructure.

Current statistical methods for tractometry often conduct independent statistical tests at each bundle segment, and thereby treat these spatial bins as independent hypotheses. BUAN and AFQ-Insight (Kruper et al., 2025) apply pointwise false discovery rate (FDR) correction using the Benjamini-Hochberg (BH-FDR) procedure (Benjamini & Hochberg, 1995). To account for the spatial autocorrelations along tracts, BUAN 2.0 models streamlines using functional data analysis, such as Function-on-Scalar (FOSR) regression (B. Chandio et al., 2023; B. Q. Chandio, 2022; Ramsay & Silverman, 1997), and AFQ’s Tractable R package (Kruper et al., 2025) integrates a first-order autoregressive model with generalized linear models (GLM). However, these approaches are primarily designed for single bundles that are modeled using 1D parameterizations. Studies investigating microstructural abnormalities, such as the ENIGMA TBSS protocol (Kelly et al., 2018; Van Velzen et al., 2020; Villalón-Reina et al., 2020), typically examine multiple ROIs from a given atlas (e.g., the JHU atlas Mori et al., 2008). Extending such studies to tractometry requires some type of multiple testing correction to be applied to all subdivisions across all bundles being studied. With 2D parameterization, the number of tests can quickly become very large, and may require more complex models of spatial correlation, to estimate dependencies among statistical tests. Effective statistical inference for multi-bundle tractometry should therefore (i) derive localized test statistics at each spatial bin for interpretability, yet (ii) still maintain error control jointly across all bins and all bundles, and (iii) account for the spatial correlation structure induced by the bundle geometry and spatial parameterization framework.

To address these requirements, we introduce *SPECTRA, Spatial Inference for Tractometry*, for advanced streamline-based 2D parameterization with spatially-aware statistical inference for multi-bundle tractometry. For inference, we adapt the two-stage hierarchical FDR (hFDR) framework of Benjamini and Heller, 2007, which first tests coarse spatial clusters before cell-level trimming within rejected clusters. We define data-adaptive clusters by estimating the empirical spatial correlation structure using a Matérn kernel (Matérn, 1986; Stein, 1999) per bundle and using the resulting length scales for partitioning, analogous to resolution elements (RESELs) in neuroimaging random field theory (Worsley et al., 1992, 1996). We evaluate SPECTRA through extensive simulation experiments by injecting synthetic effects at varying locations, spatial extents, and magnitudes into real multi-site dMRI data, with a particular emphasis on *data efficiency*, i.e., how strong the detected effects are over a range of sample sizes. Given recent interest and debate over the sample sizes required for reproducible brain-wide association studies (Marek et al., 2022; Spisak et al., 2023; Tervo-Clemmens et al., 2023), improving data efficiency through principled statistical design is of broad importance. Our framework pursues this by incorporating spatial structure into the inference procedures. We show that the hierarchical structure can boost power and data efficiency of tractometry statistics while maintaining error control, compared to pointwise BH-FDR applied across all bundles simultaneously. We then demonstrate SPECTRA by examining the effects of dementia and mild cognitive impairment (MCI) on local microstructure in a relatively large dataset of over 4,000 subjects across 63 WM bundles.

SPECTRA is available as an open-source Python package at https://github.com/wendyfyx/SPECTRA.

## 2 Methods

### 2.1 Data and Preprocessing

#### Participant Demographics

We analyzed dMRI data from 3,049 subjects in the Health & Aging Brain Study (HABS-HD) dataset (O’Bryant et al., 2021) and 1,215 subjects from Phase 3/4 of the Alzheimer’s Disease Neuroimaging Initiative (ADNI) dataset (Jack et al., 2010). A total of 4264 subjects includes 2966 cognitively normal (CN), 981 with mild cognitive impairment (MCI) and 317 with dementia. Participant demographics by scanner or protocol are summarized in Table 1.

**Table 1:** Demographic summary by dataset and protocol.

| Dataset | Protocol* | N(Total) | N(CN)<sup>†</sup> | N(MCI)<sup>†</sup> | N(Dementia) | Sex (F/M) | Age (mean ± sd) |
| --- | --- | --- | --- | --- | --- | --- | --- |
| ADNI | GE127<sup>‡</sup> | 29 | 21 | 8 | 0 | 20/9 | 68.8 ± 6.5 |
| | GE36 | 70 | 40 | 25 | 5 | 39/31 | 71.3 ± 6.8 |
| | GE54 | 166 | 94 | 50 | 22 | 78/88 | 75.0 ± 8.0 |
| | P100<sup>‡</sup> | 25 | 17 | 6 | 2 | 20/5 | 69.3 ± 6.2 |
| | P33 | 67 | 41 | 19 | 7 | 29/38 | 75.7 ± 7.5 |
| | P36 | 59 | 22 | 31 | 6 | 29/30 | 72.9 ± 6.6 |
| | S100<sup>‡</sup> | 11 | 5 | 5 | 1 | 7/4 | 70.1 ± 8.0 |
| | S127<sup>‡</sup> | 354 | 226 | 99 | 29 | 216/138 | 71.0 ± 8.4 |
| | S31 | 104 | 65 | 29 | 10 | 62/42 | 71.6 ± 8.4 |
| | S55 | 330 | 197 | 97 | 36 | 180/150 | 73.8 ± 8.0 |
| | - | 1215 | 728 | 369 | 118 | 680/535 | 72.6 ± 8.1 |
| HABS-HD | Skyra | 1724 | 1359 | 254 | 111 | 1058/666 | 66.7 ± 8.8 |
| | Vida1 | 1325 | 879 | 358 | 88 | 854/471 | 64.1 ± 8.4 |
| | - | 3049 | 2238 | 612 | 199 | 1912/1137 | 65.6 ± 8.7 |
| Total | - | 4264 | 2966 | 981 | 317 | 2592/1672 | 67.6 ± 9.1 |

\* In the ADNI DTI protocol abbreviations, scanner vendors are denoted by GE (General Electric), P (Philips), and S (Siemens), and the number denotes the total number of diffusion gradients in the protocol.
<sup>†</sup> CN: cognitively normal; MCI: mild cognitive impairment.
<sup>‡</sup> For multi-shell protocols, dMRI volumes with the lowest nonzero b-values between 600 and 1600 (tolerance=20) were extracted and used to fit DTI, and msmtCSD was used to generate fODFs instead of CSD.

Data used in the preparation of this article were obtained from the ADNI database (adni.loni.usc.edu). The ADNI was launched in 2003 as a public-private partnership, led by Principal Investigator Michael W. Weiner, MD. The primary goal of ADNI has been to test whether serial magnetic resonance imaging (MRI), positron emission tomography (PET), other biological markers, and clinical and neuropsychological assessment can be combined to measure the progression of mild cognitive impairment (MCI) and early Alzheimer’s disease (AD).

#### Diffusion MRI Preprocessing

For both datasets, dMRI data was preprocessed with the following steps: denoising (Cordero-Grande et al., 2019; Veraart et al., 2016), Gibbs ringing correction (Kellner et al., 2016; H.-H. Lee et al., 2021), susceptibility-induced distortion correction with FSL’s *topup* (Andersson et al., 2003), eddy current and motion correction with FSL’s *eddy* (Andersson & Sotiropoulos, 2016), and bias field inhomogeneity correction (Tournier et al., 2019). Preprocessing details can be found in Liou et al., 2025; Nir et al., 2013; Thomopoulos et al., 2021. For each processed dMRI, diffusion tensors were fitted using non-linear least-squares to generate scalar maps of fractional anisotropy (FA), mean (MD), radial (RD), and axial diffusivity (AxD) in the subject’s native space (Le Bihan et al., 2001). Constrained spherical deconvolution (CSD) (Tournier et al., 2007) was used to reconstruct fiber orientation distribution functions (fODFs), except in the case of the ADNI protocols, GE127, P100, S100 and S127, where multi-shell multi-tissue CSD (msmtCSD) was used (Jeurissen et al., 2014).

#### Bundle Tractography

To reconstruct bundles from processed dMRI, we used a symmetric adaptation of the HCP1065 atlas (Ba Gari et al., 2026; Yeh, 2022) to mitigate hemispheric bias. Tractography was performed separately for each bundle by constraining the tracking region using atlas bundles as references (Feng et al., 2026; Rheault et al., 2019; Yeh, 2020). Each subject’s FA, MD, and the mean b0 image were registered to the ICBM 2009a Nonlinear Asymmetric T1w template using multichannel ANTs SyN (Tustison et al., 2021). Each atlas bundle was converted to a binary mask, dilated by 5mm, and warped to subject dMRI space via the inverse transform to define the seed region. Eight seeds per voxel were placed within this mask. The stopping criterion was the intersection of the dilated seed mask with an adaptive FA threshold, defined as the 50th percentile of the subject’s FA value, lower bounded by 0.3, ensuring adequate tracking in regions of potentially reduced anisotropy. DIPY’s local probabilistic tractography(Garyfallidis et al., 2014) was performed in the subject’s native space with a step size of 0.5 mm. Streamlines were retained if their lengths were within the atlas bundle range with a 20% tolerance. Atlas bundles were then warped to subject space and used to filter streamlines with the Fast Streamline Search (FSS) algorithm (St-Onge et al., 2022, 2023). The max angle used for tracking and the FSS radius for each bundle are listed in **Supplementary section 1**. The resulting bundles were kept in the subject dMRI space and also warped to the MNI space for bundle profiling.

### 2.2 Tractometry and Spatial Parameterization

We designed the following 2D tractometry approach to quantify microstructural metrics along and across extracted WM bundles, illustrated in Figure 1. A 2D grid with along-tract (*s*) and radial (*r*) dimensions was first defined on an atlas bundle in the MNI space, then applied to subject bundles. Both 1D and 2D parameterizations are available in the SPECTRA package.

**Figure 1:**
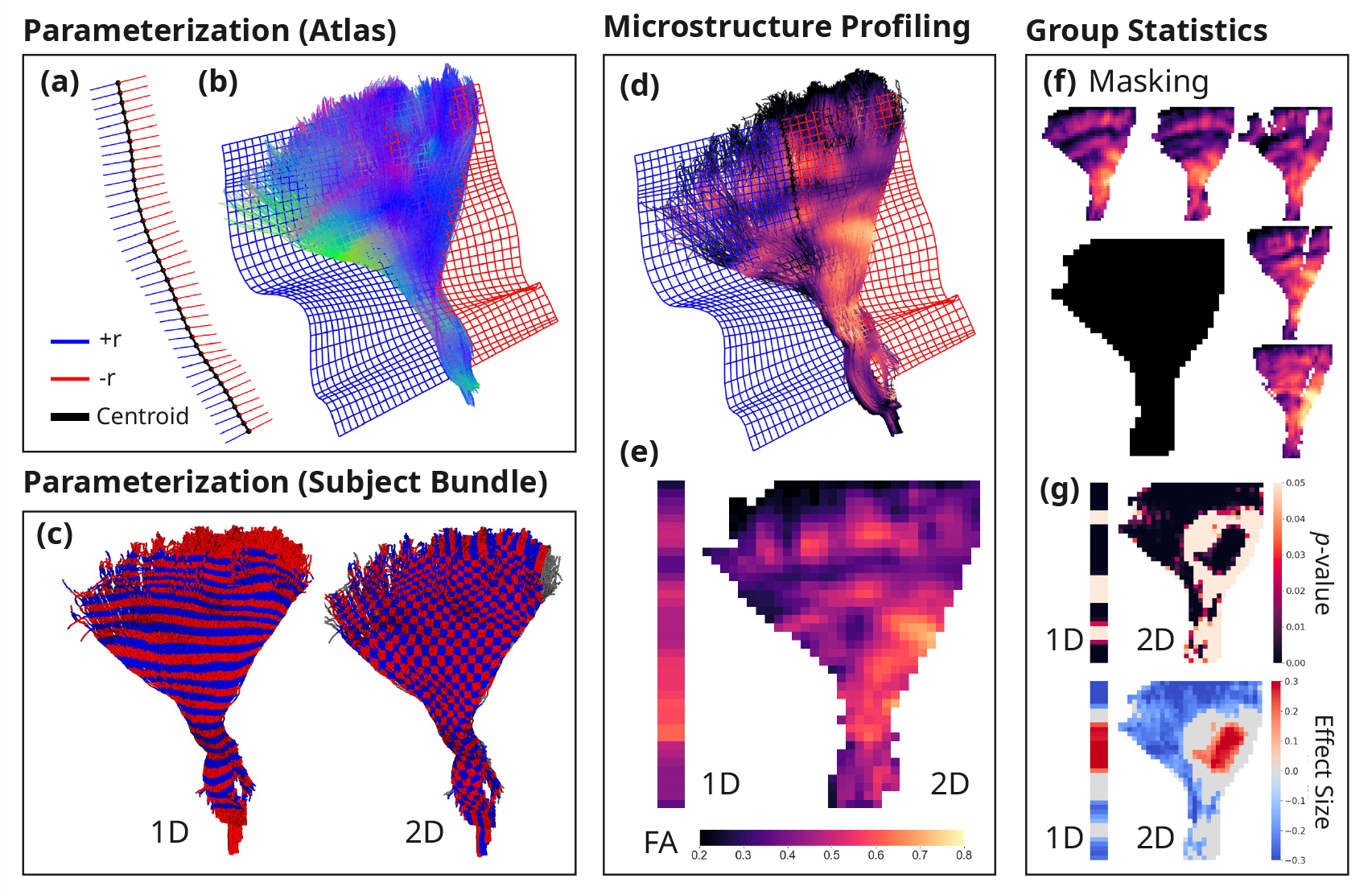
2D bundle parameterization, illustrated here with the left frontal cortico-pontine tract. **(a)** The atlas bundle centroid is used to derive fixed-sized along-tract segments, and the radial vectors are defined as the direction with the maximal variance for each segment using PCA. **(b)** Fixed-sized radial bins are computed from the signed radial distances for streamline points on the atlas bundle. Together with along-tract segments, a 2D grid is defined using the atlas bundle. **(c)** An example subject bundle is parameterized with both 1D and 2D tractometry using the grid computed in step (b). **(d)** A scalar FA map projected to all streamline points from the subject bundle in the native space. **(e)** Projected FA values are averaged per grid cell to create the 2D profile. The 1D FA profile is shown for comparison. **(f)** Subjects’ FA profiles are aggregated to create a group-level mask, where grid cells that have data from less than 30% of the subjects are removed. **(g)** Linear mixed model is fitted at each grid cell to obtain a spatial map of p-values and effect size. The same procedure is performed for each segment from 1D profiles for comparison.

#### Along-Tract Parameterization

Along-tract bins (*s*-bins) were defined using BUAN’s segment assignment method (B. Q. Chandio et al., 2020). The atlas bundle centroid derived from QuickBundles (QB) (Garyfallidis et al., 2012) was resampled to 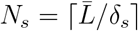 equidistant points, where 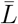 is the mean streamline length and *δ*_*s*_ is the desired segment length (mm). Each streamline point from the atlas bundle was assigned to its nearest centroid point using a KD-Tree, defining the *s*-bin membership. Terminal bins containing fewer than 20% of the median bin point count were merged with their neighbors.

#### Robust Centroid Extension

When a bundle contains streamlines with highly variable lengths, the QB centroid may be shorter than the full bundle extent, causing the terminal *s*-bins to span disproportionately large regions. To address this, we optionally extend the centroid to match the true bundle extent, see Figure 2(a). The QB centroid is extended by 30% of its arc length, by extrapolating linearly from each endpoint using the terminal tangent vectors. The final centroid extent is then determined by projecting all streamline endpoints onto the extended curve via KD-Tree, and taking the 2nd and 98th percentiles of the projected positions. The clipped portion is resampled to *N*_*s*_ equidistant points via arc-length interpolation.

**Figure 2:**
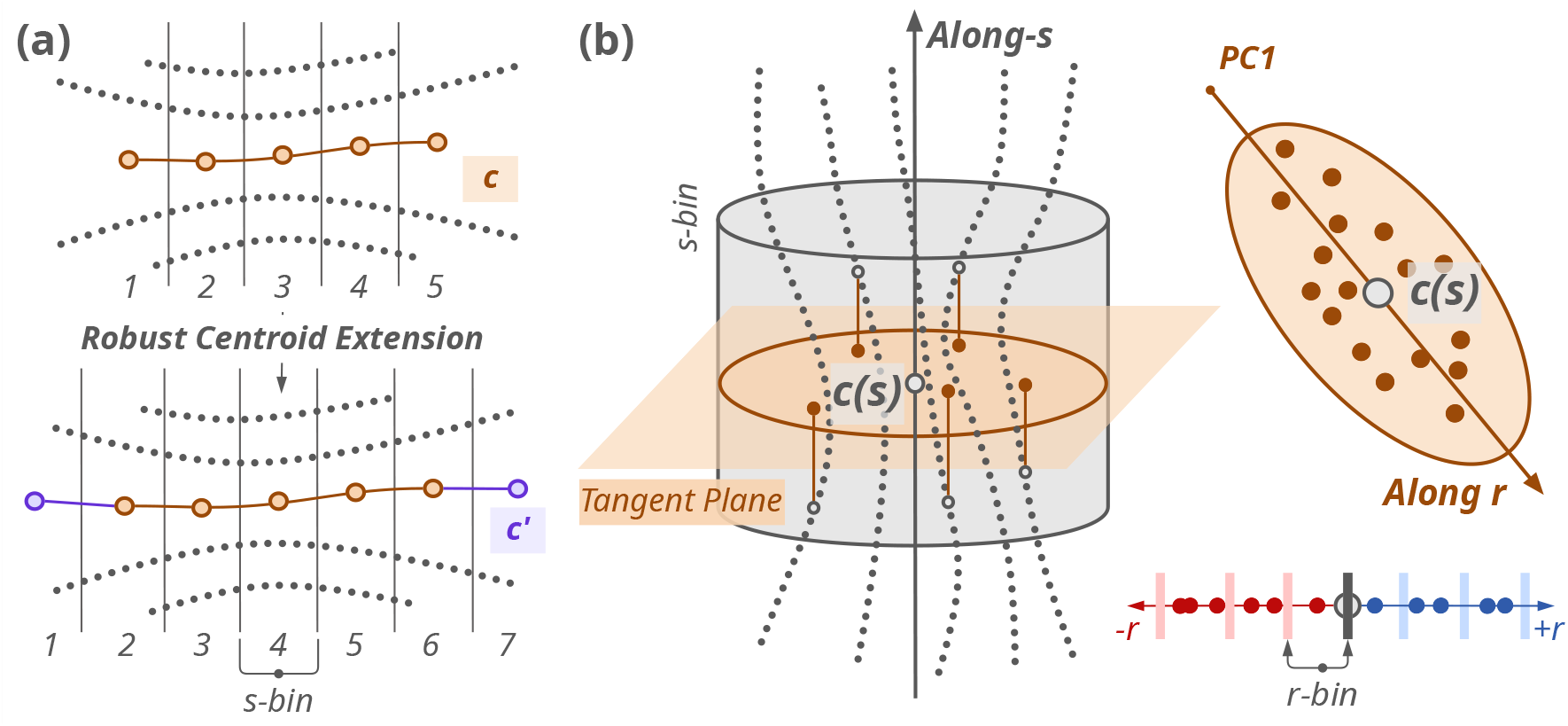
Illustrative example of the (a) robust centroid extension used for creating along-tract segments and (b) radial parameterization. *c*: centroid; *c*^*′*^: extended centroid; *c*(*s*) centroid point for segment *s*.

#### Radial Parameterization

At each centroid point *c*(*s*), we defined a local tangent plane perpendicular to the centroid trajectory. All streamline points assigned to bin *s* were projected onto this tangent plane, and principal components analysis (PCA) was applied to the projections. The first principal component defined the radial unit vector ***u***(*s*), representing the direction of maximum variance, see Figure 2(b). To ensure continuity along the tract, radial vectors were oriented consistently along consecutive segments and smoothed using a 1D Gaussian filter. Signed radial distances were then computed for each streamline point along ***u***(*s*) and binned into *N*_*r*_ *r*-bins of width *δ*_*r*_ (mm). Edge bins containing fewer than 20% of the median bin count were merged with their neighbors, yielding a fixed (*N*_*s*_, *N*_*r*_) grid for the atlas bundle.

#### Subject Bundle Parameterization

Subject bundles registered to the MNI space were parameterized using the same grid defined on the atlas bundle. Each streamline point on the subject bundle was assigned to its nearest atlas centroid (*s*-bin), and signed radial distances were computed using the atlas radial vectors ***u***(*s*) and binned using the atlas bin edges. Two validity checks were then applied to ensure that we do not make inferences in regions with poor coverage at the subject and group level. First, streamline points beyond the atlas grid’s radial extent were excluded. Second, grid cells with fewer than a threshold number of streamline points were masked as invalid. We defined the threshold as 20% of the median point count across grid cells, clamped to a range between 50 and 200 points. This prevents overly lenient masking in high-density bundles and overly strict masking in low density ones.

#### Microstructure Profiling

For each metric analyzed, the microstructural scalar map (e.g., FA) was sampled at each streamline point in the native space bundle, and averaged within each valid grid cell, producing a subject-specific 2D profile of size (*N*_*s*_, *N*_*r*_). For 1D profiling, BUAN provides two complementary bundle profiling approaches: (i) a full-resolution strategy that utilizes all streamlines and all points within each segment, preserving intra-segment spatial detail, where multiple points within a segment are accounted for by modeling them as random effects in linear mixed models (B. Q. Chandio et al., 2020), and (ii) a compact representation that summarizes each segment using a 1D weighted mean profile (B. Q. Chandio et al., 2024a). In this study, we used BUAN’s weighted mean approach to create the 1D profiles, where point-wise measures are weighted by their distance to the atlas centroid *s*-bin point and averaged within each segment. After individual subject parameterization, a group-level validity mask was constructed, retaining only cells with valid data in at least 30% of subjects. The 2D parameterization pipeline is summarized in Figure 1. Bundle profiles of FA, MD, RD and AxD were extracted in both 1D and 2D parameterization at a spatial resolution of 5 mm along-tract and 5 mm radially, for 63 WM bundles (see **Supplementary section 1**. All subjects with complete demographic data and valid profiles for at least one bundle were retained.

### 2.3 Statistical Modeling

At each spatial bin (*s, r*) of each bundle, we fit the following linear mixed model (LMM):

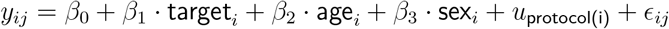

where *y*_*ij*_ is the microstructural metric (e.g., FA) for subject *i* at bin *j, β*_0_ is the intercept, *β*_1..3_ are the fixed effects, *u*_site(i)_ is the random intercept accounting for variability across MRI protocols, *ϵ*_*ij*_ ∈ 𝒩(0, *σ*^2^) represents the residual error. The target variable may be categorical (e.g., diagnosis) or continuous (e.g., cognitive score). Continuous outcomes and fixed effect predictors were standardized across all spatial bins per bundle prior to model fitting. Models were fit using the *lmer* function via *pymer4* (Jolly, 2018). For categorical targets, we also compute Cohen’s d as a standardized effect size measure.

### 2.4 False Discovery Rate (FDR) Correction

Analyzing a large number of spatial bins for 2D parameterized bundles requires principled control of false discoveries. We implement three FDR correction strategies in SPECTRA, described in the following sections, progressing from standard global correction to a spatially-informed hierarchical procedure.

#### 2.4.1 The Benjamini-Hochberg Procedure

Let *p*_(1)_ ≤ *p*_(2)_ ≤ … ≤ *p*_(*M*)_ denote the ordered raw *p*-values across all *M* valid bins pooled across all bundles. The Benjamini-Hochberg (BH) procedure (Benjamini & Hochberg, 1995) rejects hypothesis *H*_(*j*)_ if

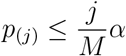

where *α* is the target FDR level. Equivalently, BH-adjusted *p*-values (*q*-values) are defined as

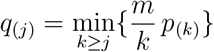

and a bin is declared significant if *q*_(*j*)_ *< α*.

#### 2.4.2 Adaptive FDR with Storey’s Estimator

To account for the possibility that many hypotheses are truly non-null, we also implement an adaptive variant of the global FDR procedure using Storey’s *π*_0_ estimator (Storey, 2002). The BH procedure assumes that all *M* tests are null, i.e. *π*_0_ = 1, whereas Storey’s estimator provides a data-driven estimate of 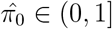. A cubic spline is fitted to

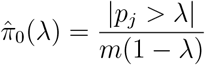

across a grid *λ* ∈ [0.05, 0.95] of threshold values, and extrapolated to *λ* = 1 to yield 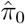. The effective FDR threshold becomes 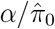, increasing power when many hypotheses are non-null. A safeguard is applied so that the *q*-values are lower-bounded by their corresponding raw *p*-values, preventing paradoxical rejections when 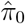 is very small (Reiss et al., 2012).

#### 2.4.3 Spatially-Informed Hierarchical FDR

The global BH procedure and the adaptive variant treat all bins from bundles as independent. SPECTRA accounts for the spatial structure of WM bundles by assuming that residuals from the LMM are likely to exhibit smooth spatial correlation. We propose the following inference procedure by first estimating the spatial autocorrelation via a Matérn kernel, and using it to define blocks for inference. We adopt the hierarchical FDR (hFDR) framework of Benjamini and Heller, 2007, which controls FDR across two levels of a spatial hierarchy. This procedure operates in two stages: a coarse block-level testing (Stage 1) followed by a cell-level trimming within the significant blocks (Stage 2).

#### Estimating Spatial Correlation via Matérn Kernel

A Matérn covariance function is fitted to the empirical autocorrelation *ρ*(*m*) = *C*(*m*)*/C*(0) at lag *m* of the reduced-model residuals, along both axes of each bundle’s grid. Using the reduced-model residuals ensures that the estimated correlation structure reflects noise rather than the effect of the target variable. The Matérn family is parameterized by a smoothness parameter *ν*:

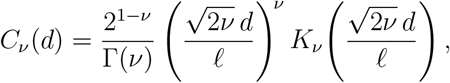

where *d* is the physical separation between bins (mm), *𝓁* is the fitted length scale (mm), *K*_*ν*_ is a modified Bessel function, and Γ is the gamma function. At *ν* = 1*/*2 this reduces to the exponential kernel *C*(*d*) = exp(−*d/𝓁*), while as *ν* → ∞ it converges to the radial basis function (RBF) kernel *C*(*d*) = exp(−*d*^2^*/*2*𝓁*^2^). The length scales *𝓁*_*s*_ and *𝓁*_*r*_ are fitted separately along each axis, and the full 2D covariance between bins (*s*_*i*_, *r*_*i*_) and (*s*_*j*_, *r*_*j*_) is assumed to be separable:

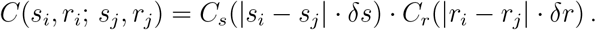

Length scales *𝓁*_*s*_ and *𝓁*_*r*_ are estimated by minimizing the weighted sum of squared residuals between the empirical and modeled autocorrelation, with each lag weighted by its number of contributing observation pairs. Fitting uses lags *m* ≥ 1 up to the first lag at which 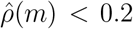, to exclude noisy tails. The *ν* = 1*/*2 kernel consistently achieves the best fit across the bundles and DTI metrics examined, as illustrated in Figure 3(a), and is therefore the default in SPECTRA and used in subsequent experiments.

**Figure 3:**
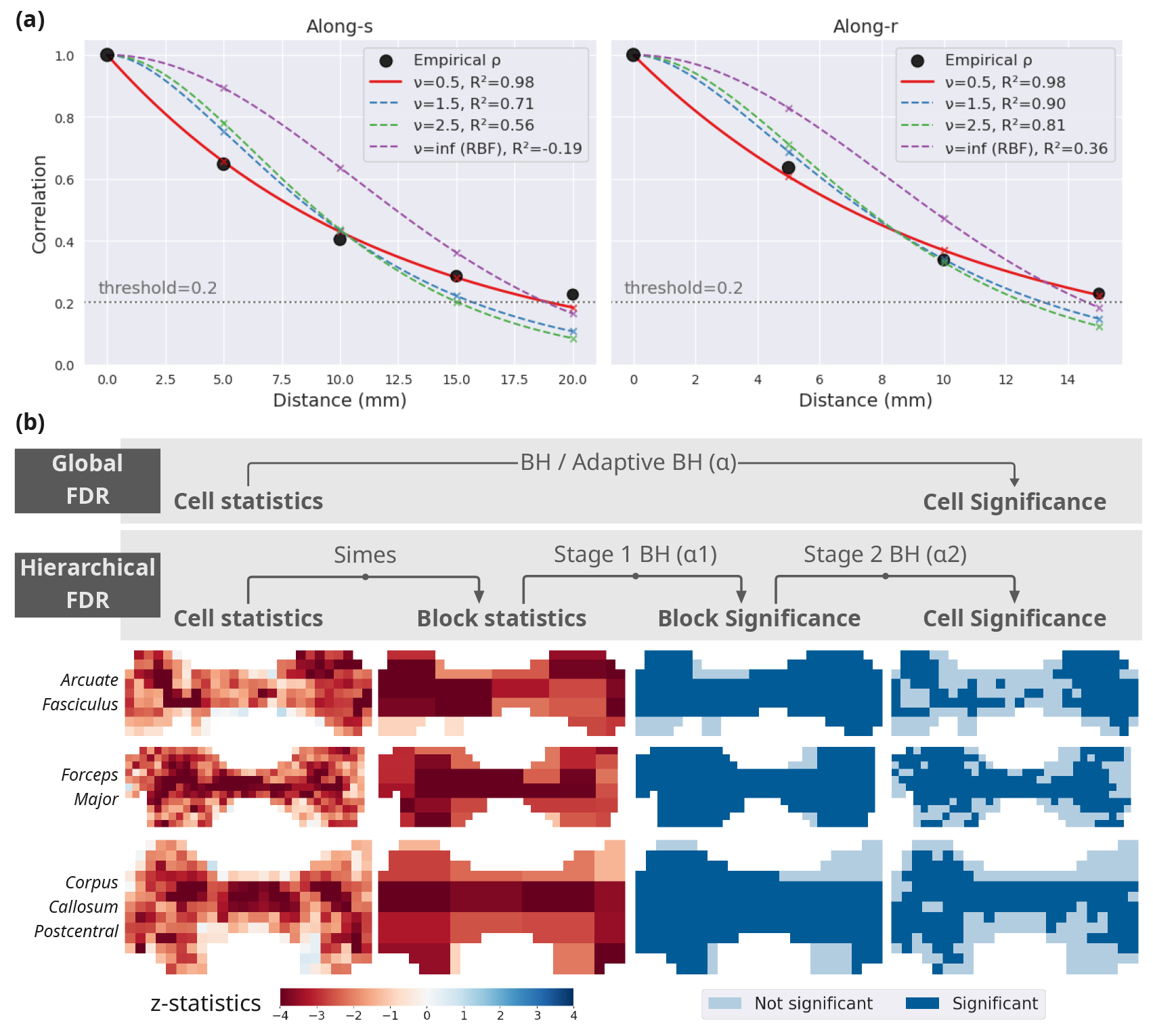
**(a)** Example of Matérn kernel fit across four values of *ν* along *s*− and *r*− axes of FA profiles for the corpus callosum postcentral bundle. *R*^2^ was used to evaluate model fit. The empirical correlation *ρ* measures at different values of *d* are shown as black dots. **(b)** Hierarchical FDR (hFDR) with RESEL blocks, where the block sizes were determined from the fitted Matérn kernels. In hFDR-RESEL, the BH procedure was applied on block-level Simes statistics in Stage 1 and the conditional p-values in Stage 2, to derive the final cell-level significance, as opposed to the global FDR procedure which applies to the cell-level statistics only. Note that all BH procedures were applied to statistics aggregated across all bundles in the analysis.

Analogous to the resolution element (RESEL) concept in random field theory (Worsley et al., 1996), we define the RESEL density (Worsley, 2002) as a measure of the number of effective resolution elements per unit area after accounting for spatial correlation. In random field theory, RESELs are dimensionless quantities derived from the ratio of the search region volume to the smoothness of the field, typically parameterized by the full-width at half maximum (FWHM) of the residual field. The total RESEL count reflects the effective number of resolution elements contributing to the topology of statistical maps, and governs the expected Euler characteristic used for multiple comparisons correction.

The total RESEL count approximates the number of independent observations in the volume. Here, the FWHM along-tract dimension is derived from the fitted Matérn length scale as FWHM = 2.355 *𝓁*, and the RESEL density is defined as the ratio of the grid cell area to the FWHM area:

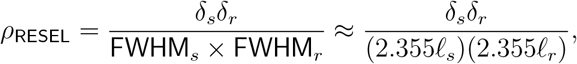

and *ρ*_RESEL_ ≈ *δ*_*s*_*/*(2.355*𝓁*_*s*_) for 1D parameterized grids. The RESEL density is used here solely to define block sizes for the hierarchical FDR procedure described below.

##### Blocking Partitioning

For stage 1 testing in the hFDR procedure, each bundle is partitioned into non-overlapping rectangular blocks whose lengths along *s* and *r* are set to one FWHM (2.355*𝓁*_*s*_*/δ*_*s*_ and 2.355*𝓁*_*r*_*/δ*_*r*_ bins respectively). This ensures that cells within a block are strongly correlated while cells in different blocks are approximately independent. Blocks with fewer than 3 cells are merged to their nearest neighbor. As a simple alternative to RESEL-based blocking, we also evaluate a variant in which each bundle is treated as a single block. This approach is less sensitive to the accuracy of the length scale estimates and provides a baseline for comparison with RESEL blocks. We will refer to this variant as *hFDR-Bundle* mode as opposed to *hFDR-RESEL* mode.

##### hFDR Stage 1: Block-Level Testing

Within each block *b* containing *c*_*b*_ cells, let 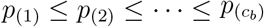 denote the ordered cell-level raw *p*-values. We compute a combined *p*-value using Simes’ test (Simes, 1986)

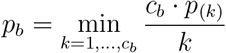

Simes’ test is valid under the assumption of positive regression dependence on a subset (PRDS), which holds for spatially correlated Gaussian statistics, and is naturally powerful for spatially localized effects because it is insensitive to dilution by null cells in a block. The BH procedure is applied across all block-level *p*_*b*_ values at level *α*_1_, see Figure 3(b). Let *u*_1_ be the largest raw *p*_*b*_ among all rejected blocks, and this serves as the Stage 1 selection threshold for Stage 2.

##### hFDR Stage 2: Cell-level Trimming

Following Stage 1 block-level testing, cell-level significance is assessed using the conditional rather than the marginal *p*-values to properly account for cells being in a rejected block (Benjamini & Heller, 2007). For cell *i* in rejected block *b*, the conditional *p* is defined as

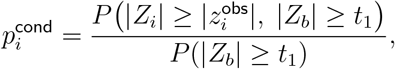

where 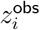 is the observed cell *z*-score, *Z*_*b*_ is the block-level test statistic, and *t*_1_ = Φ^−1^(1 − *u*_1_*/*2) is the two-sided *z*-threshold corresponding to *u*_1_. Following Benjamini and Heller (2007), we use the conservative upper bound on the true conditional *p*-values evaluated under the global null, whereas a cell with true signal would be more likely to appear in a rejected block, yielding a smaller conditional *p*-value.

To evaluate this numerically, we model the joint distribution of *Z*_*i*_ and *Z*_*b*_ as bivariate normal with correlation *ρ*_*i*_. Because Simes’ test does not produce a closed-form expression for the correlation between an individual cell statistic and the block statistic, we assume equal and independent contributions from all cells, giving 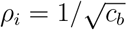. The conditional *p*-value then evaluates to:

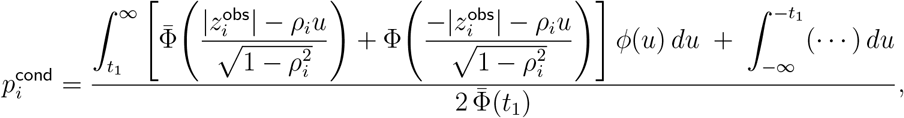

where *ϕ* and Φ are the standard normal PDF and CDF, 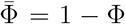, and the integrand in the second term is identical to that in the first. The BH procedure is then applied to 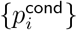 within each rejected block at *α*_2_ to produce cell-level significance, see Figure 3(b).

### 2.5 Simulation Experiments

To evaluate and compare how statistical inference methods can detect effects of different magnitudes, spatial extent and geometry across different sample sizes, we designed the following simulation experiments by injecting synthetic effects into real 2D tractometry profiles from control subjects. This preserves the empirical covariance structure of the data while permitting evaluation of known effects.

#### Effect Simulation

For each simulation repetition, 2*N*_*S*_ subjects were randomly sampled without replacement from the full subject pool of 2,966 CN subjects and divided into two groups of equal size *N*_*S*_. The first group received effects and the second served as controls. We also randomly selected *B* bundles to receive the synthetic effects, and for each affected bundle *b*, a spatial effect map *e*_*b*_(*s, r*) ∈ [0, 1] was constructed. One or more elliptical blobs were generated with a peak amplitude of *d*_max_ at the blob center and cosine decay toward the boundary. A minimum effect floor of *d* = 0.1 was applied to ensure that the effect map was non-trivially non-zero. We simulated two spatial configurations of effect geometry to represent two biologically plausible regimes (see Figure 4): in *diffuse* mode, a single elliptical blob was generated, occupying 30 − 50% of the grid in each dimension, producing a broad signal spanning a large tract region; in *localized* mode, four small elliptical blobs, each covering 5 − 15% of the grid, were placed at randomly locations producing spatially concentrated signals, to represent focal alterations confined to small patches. Synthetic effects were injected additively into the FA profiles of each case subject, scaled by the local standard deviation across subjects. A linear mixed model (LMM) was fitted at each grid cell following the procedure described in subsection 2.3, with the synthetic effect as the contrast.

**Figure 4:**
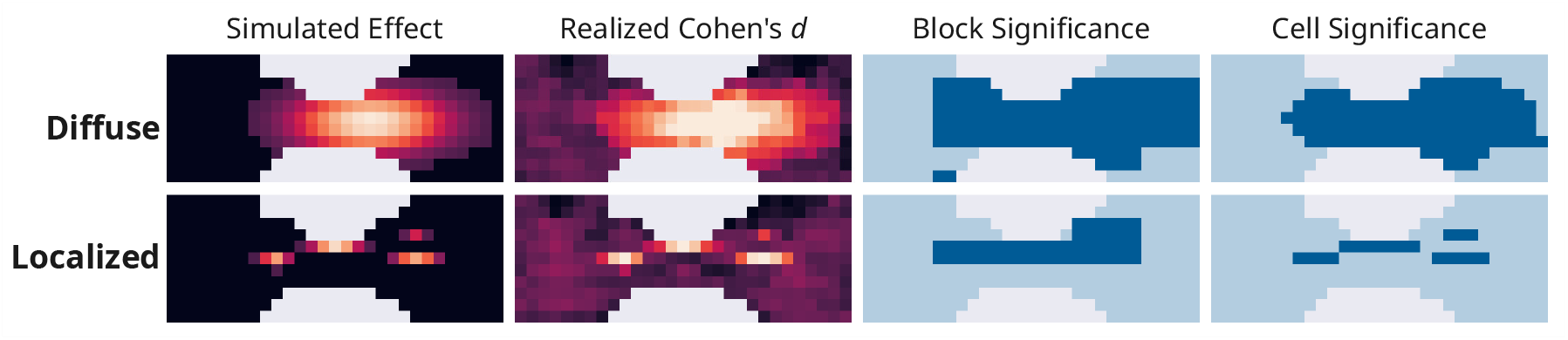
Examples of injected synthetic effects, shown on 2D grids of the *corpus callosum forceps major* (brighter colors indicate larger effect sizes). Cell-significance maps derived from FDR corrections, and additional block-significance maps for hFDR-RESEL, were evaluated against the simulated effect maps (dark blue: significant; light blue: not significant).

#### Experimental Conditions

We used a full factorial design across four factors, yielding 144 unique conditions (Table 2). Each condition was repeated for 100 repetitions with a different random seed to generate different subject pools and effect maps. For efficiency, experiments were restricted to a representative subset of 20 bundles (see **Supplementary section 1**). Four correction methods were evaluated at each repetition using identical LMM fits, so that any differences in results reflect the correction procedure alone:

**Table 2:**
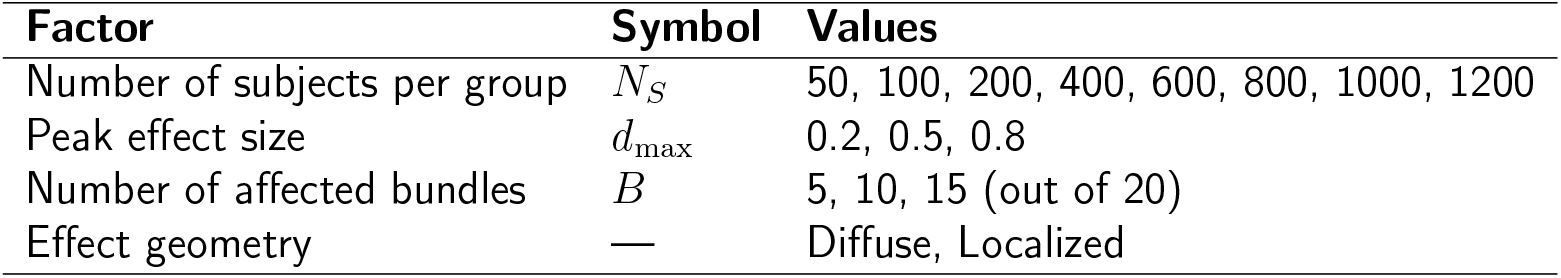
Simulation experiment factors and levels.

1. **BH-FDR**: the standard BH procedure at *α* = 0.05 applied globally across all bins from all bundles.
2. **Adaptive BH-FDR**: Storey’s 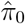-adjusted BH at *α* = 0.05 applied globally across all bins from all bundles.
3. **hFDR-Bundle**: hierarchical FDR with each bundle as a single block (*α*_1_ = 0.05, *α*_2_ ∈ {0.15, 0.25}).
4. **hFDR-RESEL**: hierarchical FDR with RESEL blocks (*α*_1_ = 0.05, *α*_2_ ∈ {0.15, 0.25}).

#### Evaluation

Performance was evaluated at two levels: At the cell level, a grid cell was labeled as a true positive if its location received a non-zero simulated effect *e*_*b*_(*s, r*) *>* 0. At the block level, a block was labeled as a true positive if any of its constituent cells received a non-zero simulated effect. We computed *sensitivity* (TP*/*(TP + FN)) and the *false discovery proportion* (FDP; FP*/*(FP + TP)).

#### Sample Size Requirements

A method is considered more *data efficient* when its sensitivity curve rises more steeply at the origin, indicating that fewer subjects are required to reach a given signal detection level. To quantitatively summarize the sample size requirements for each method, we fit a Gompertz curve to the cell-level sensitivity-vs-*N*_*s*_ relationship observed across 100 repetitions for each experimental condition. The Gompertz model, *ŝ* (*N*_*S*_) = *L*+(*U* −*L*) exp(− exp(−*k*(*N*_*S*_ −*N*_0_))), captures an asymmetric S-shaped growth which is typical for power curves. Four parameters —floor (*L*), ceiling (*U*), growth rate (*k*), and inflection point (*N*_0_) were estimated with nonlinear least squares using all 100 repetitions pooled across sample size levels. From the fitted curve, we derived *N*_70_, the minimum sample size per group required to achieve 70% sensitivity, by analytically inverting the Gompertz function as *N*_70_ = *N*_0_ − log(− log((0.7 − *L*)*/*(*U* − *L*))) */ k* and rounding up to the nearest integer.

### 2.6 Empirical Experiments

We applied SPECTRA to characterize WM microstructural differences associated with cognitive impairment. LMMs were fit on both 1D and 2D bundle profiles of FA, MD, RD and AxD across all 63 bundles at each grid cell, following the procedure described in subsection 2.3. Two group contrasts were tested: Dementia (*N* = 317) vs. CN (*N* = 2, 966) and MCI (*N* = 981) vs. CN (*N* = 2, 966), with age and sex as fixed-effect covariates and acquisition protocol as a random effect. Two correction methods were applied to each set of fitted models: global BH-FDR was applied at *α* = 0.05 across all grid cells from all bundles; hFDR-RESEL was applied with *α*_1_ = 0.05 at Stage 1 and *α*_2_ ∈ {0.15, 0.25} at Stage 2. Both *α*_2_ values were evaluated by re-thresholding the conditional *p*-values without refitting any models. For each correction method, we report the global detection rate as the proportion of statistically significant grid cells. For a subset of bundles, we visualize spatial patterns of abnormalities by projecting Cohen’s *d* at significant cells onto the atlas bundle, with individual streamline points colored according to their (*s, r*) grid cell assignment. SPECTRA provides a command-line interface for visualizing statistics derived from both 1D and 2D parameterizations projected onto atlas streamlines, built with FURY (Feng et al., 2024; Garyfallidis et al., 2021).

Full cell-, block-, and bundle-level results for all correction methods are provided in the **Supplementary Tables S1-3**, including raw and corrected *p*-values, *β* coefficients, standard errors, Cohen’s *d*, and for hFDR-RESEL, Stage 1 block statistics and estimated Matérn length scales (*𝓁*_*s*_, *𝓁*_*r*_).

## 3 Results

### 3.1 Synthetic Effects Evaluation

To evaluate how different statistical correction methods recover synthetic effects, we report cell- and block-level sensitivity and FDP with relation to sample size per group (*N*_*s*_) across all experimental conditions. Results below correspond to those from 2D tractometry profiles; the analogous evaluation on 1D profiles, generated under the same experimental conditions on the corresponding 1D grids, is reported in **Supplementary section 2**.

#### Block-Level Evaluation for hFDR

Block-level sensitivity and FDP for the two hFDR variants are shown in Figure 5. As expected, sensitivity for both methods improves monotonically with sample size and benefits from a large effect magnitude. hFDR-Bundle consistently achieves higher block-level sensitivity than hFDR-RESEL across all conditions: with moderate to large effects (*d*_max_ ≥ 0.5), hFDR-Bundle reaches near-perfect block recall at relatively modest sample sizes. hFDR-RESEL is less sensitive at the block level but maintains tighter error control. FDP for both methods remains below the nominal *α*_1_ = 0.05 for most conditions, with modest inflation only at the smallest sample size tested (*N*_*s*_ = 50). Both methods can more sensitively detect diffuse effects compared to localized effects, and this gap is most pronounced for small effect sizes (*d*_max_ = 0.2).

**Figure 5:**
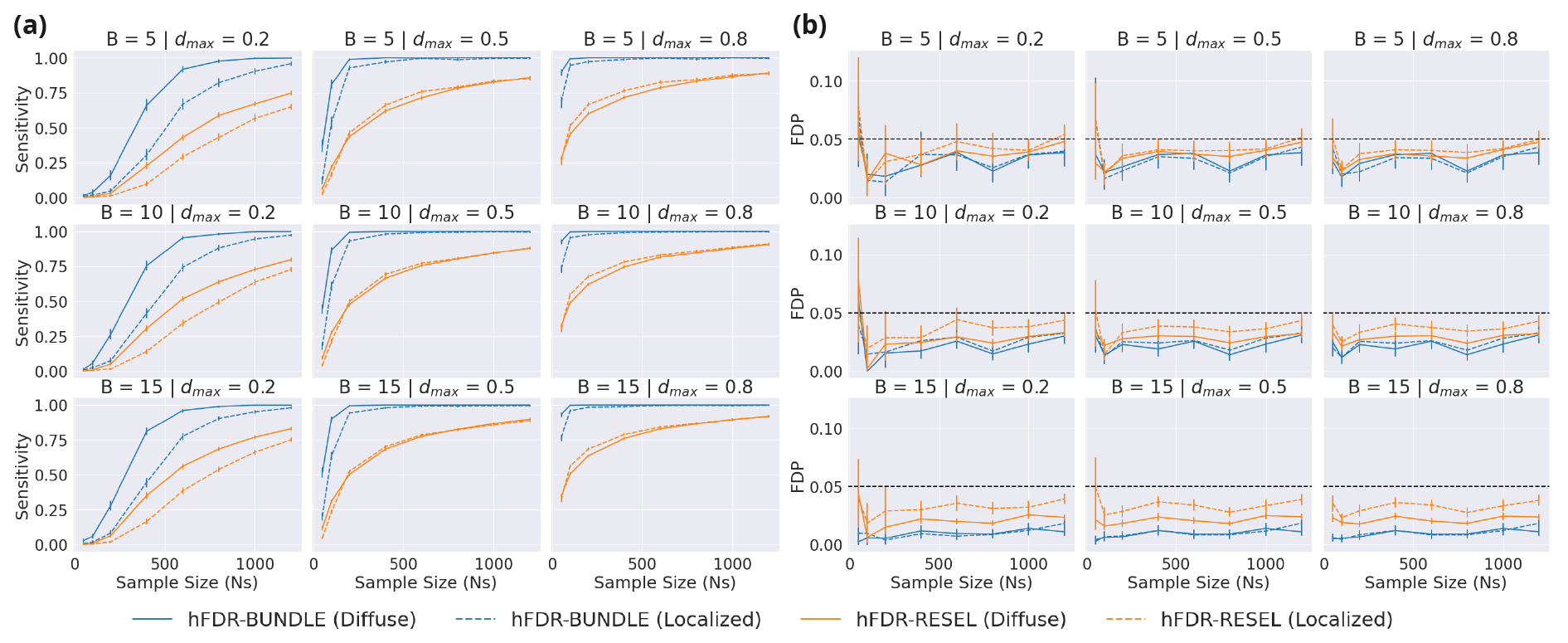
Block-level evaluation for simulation experiments for two variants of hFDR. Sensitivity (a) and FDP (b) for each experimental condition is shown with increasing sample sizes *N*_*s*_ across different peak effect sizes (*d*_max_) and numbers of affected bundles (*B*). The nominal *α* = 0.05 level is shown in panel (b) as a black dashed line.

#### Cell-Level Evaluation

Cell-level sensitivity and FDP across all four correction methods are shown in Figure 6 for diffuse (panels **(a)-(b)**) and localized (panels **(c)-(d)**) effects. All methods show better sensitivity with larger *N*_*s*_, larger *d*_max_, more affected bundles *B*, and all methods detect diffuse effects more readily than localized effects under matched conditions. Among the global FDR procedures, adaptive BH-FDR achieves modestly higher sensitivity than standard BH-FDR in diffuse settings by estimating the proportion of true nulls 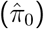, though the two methods are nearly indistinguishable for localized effects. Both hFDR variants outperform the global BH procedures at every condition, with hFDR-Bundle achieving the highest sensitivity overall. The advantage of hFDR-Bundle over hFDR-RESEL is most prominent for diffuse effects combined with low effect sizes or few affected bundles. Using a more permissive Stage 2 threshold (*α*_2_ = 0.25 vs. *α*_2_ = 0.15) improves sensitivity for both hFDR variants, and even at the more conservative *α*_2_ = 0.15, both hFDR methods retain a clear sensitivity advantage over the global procedures.

**Figure 6:**
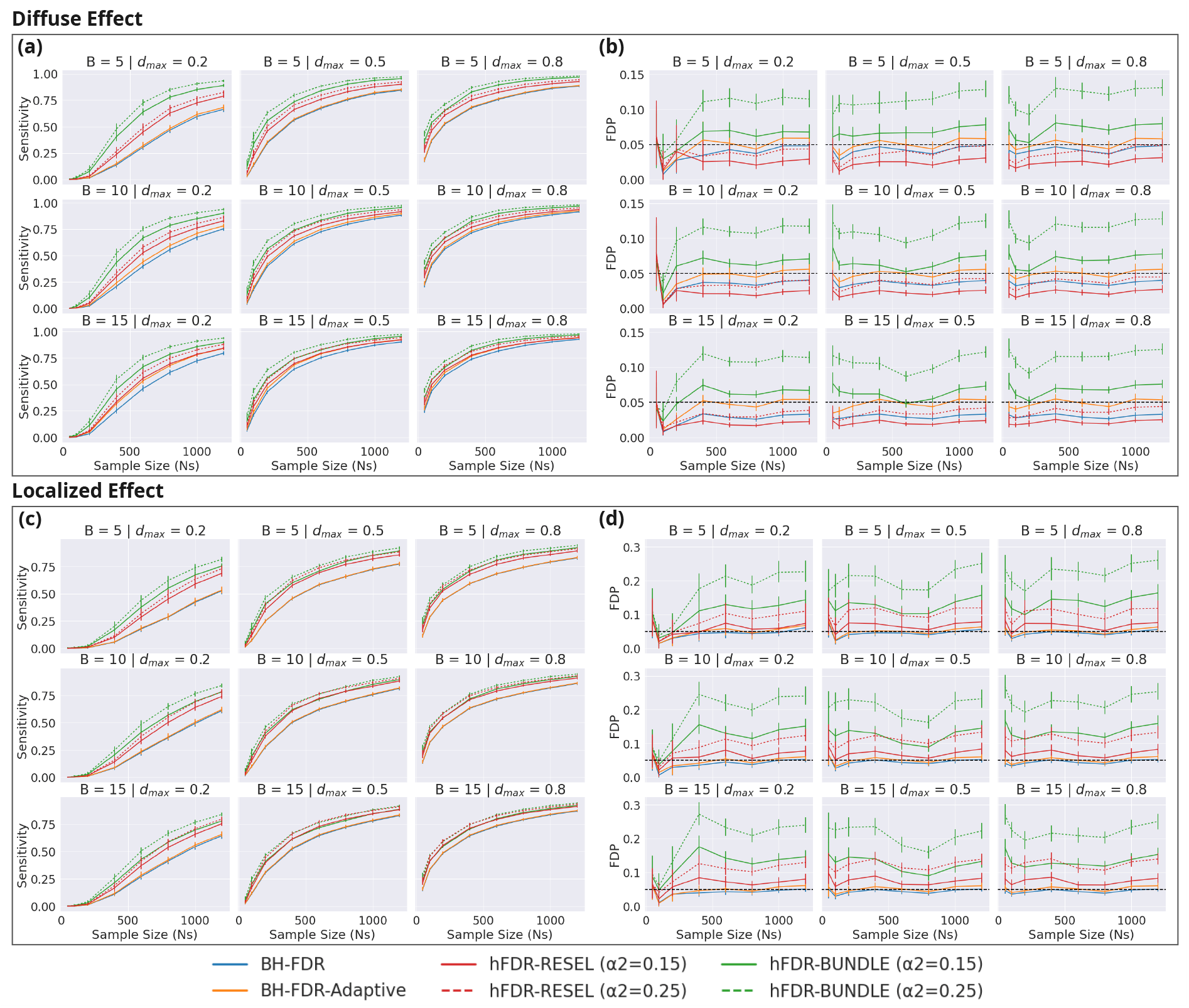
Cell-level evaluation for simulation experiments across 4 correction procedures, with different *α*_2_ shown as different line styles for the hFDR variants. Sensitivity (a, c) and FDP (b, d) for each experimental condition is plotted with increasing sample sizes *N*_*s*_ across different peak effect sizes (*d*_max_) and numbers of affected bundles (*B*), separately for diffuse and localized effects. The nominal *α* = 0.05 level is shown in panel (b, d) as a horizontal black dashed line.

All methods control the FDP at or below the nominal *α* = 0.05 for diffuse effects, except hFDR-Bundle, where FDP is inflated to 0.1-0.2 when *α*_2_ = 0.25. hFDR-RESEL at both *α*_2_ levels matches or exceeds global BH-FDR in error control. For localized effects, FDP is uniformly elevated relative to diffuse effects, consistent with the increasing difficulty and precision required to localize small signals. Under localized effects, hFDR-Bundle at *α*_2_ = 0.25 reaches FDP of 0.15-0.25, substantially exceeding the nominal level. FDP for hFDR-RESEL at *α*_2_ = 0.25 is also moderately inflated (0.1-0.15) but substantially lower than hFDR-Bundle. Reducing *α*_2_ to 0.15 brings hFDR-RESEL’s FDP to near-nominal levels even under localized effects, while hFDR-Bundle remains elevated. Taken together, these results indicate that hFDR-RESEL at *α*_2_ = 0.25 strikes a favorable sensitivity-specificity balance for diffuse spatial signals, and that lower *α*_2_ provides an additional lever for better error control when signals are more localized.

#### Sample Size Requirements

Figure 7 summarizes data efficiency via *N*_70_, the estimated sample size per group required to achieve 70% cell-level sensitivity, derived from Gompertz curve fits. Across all conditions, both hFDR variants require fewer subjects than the global BH procedures to reach the 70% sensitivity threshold. hFDR-Bundle achieves the lowest *N*_70_ values in nearly every condition: at *α*_2_ = 0.25, it requires on average only ∼51% of the subjects needed by BH-FDR for diffuse effects, and ∼58% for localized effects. hFDR-RESEL requires modestly more subjects than hFDR-Bundle but still offers a meaningful reduction, requiring approximately 65% of BH-FDR’s *N*_70_ when *α*_2_ = 0.25 and 75% when *α*_2_ = 0.15. Adaptive BH-FDR provides only a marginal improvement over standard BH-FDR, requiring ∼95% the subjects. Across all methods, *N*_70_ decreases with increasing *d*_max_ and *B*, as expected, and all methods require more subjects for localized versus diffuse effects. hFDR-RESEL is the *least* sensitive to effect geometry, with its *N*_70_ ratio between diffuse and localized conditions averages to 0.76, whereas hFDR-Bundle is the most sensitive (ratio=0.60). These results suggest that for well-powered studies targeting diffuse alterations, hFDR-RESEL offers a practically meaningful reduction in required sample size relative to global FDR correction while maintaining acceptable error control.

**Figure 7:**
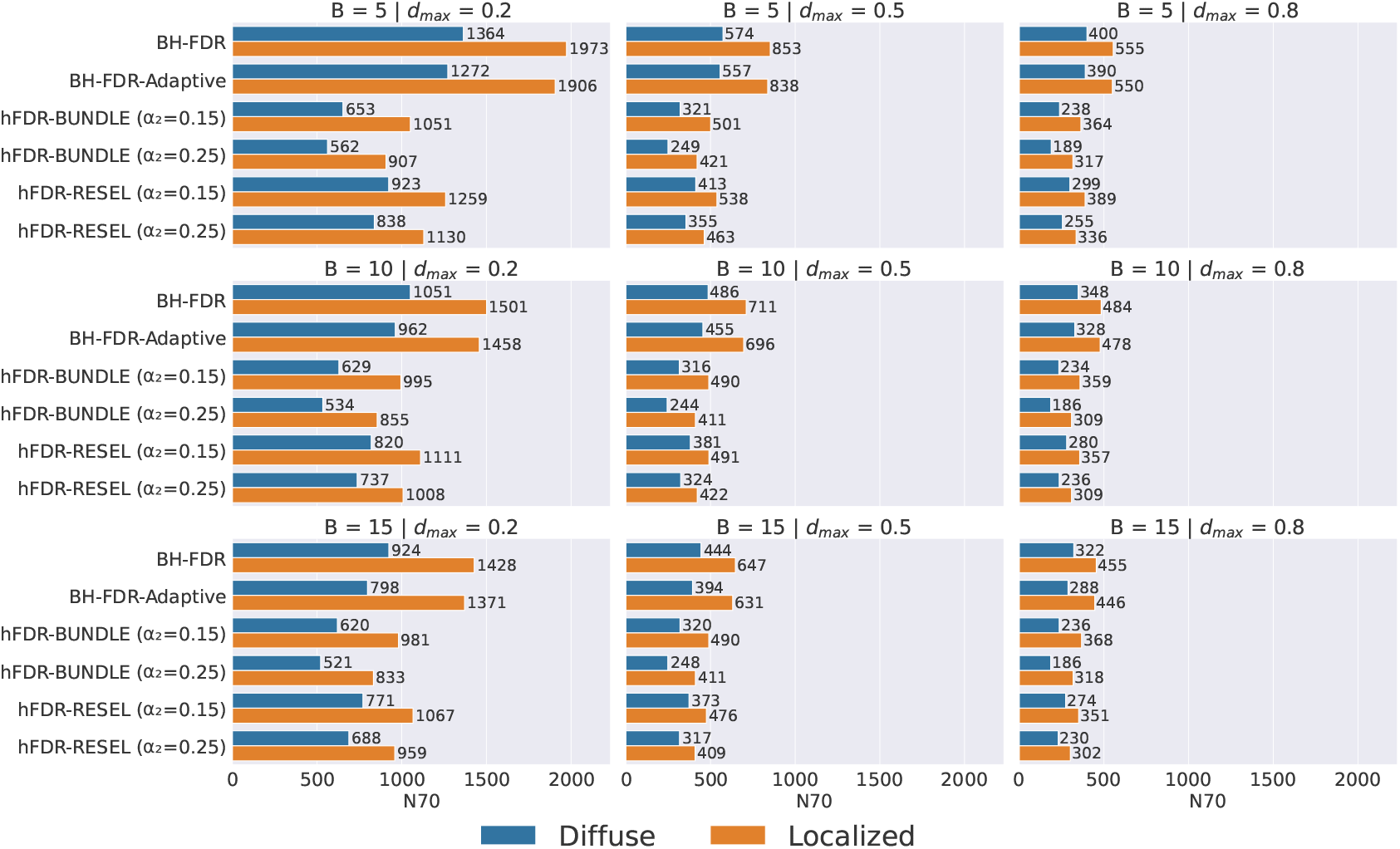
Sample size per group required to reach 70% sensitivity (*N*_70_) across various experiment conditions and correction methods, derived from fitted Gompertz curves.

### 3.2 Experimental Results on Non-Simulated Data

To evaluate SPECTRA on real data, we applied the full parameterization and statistical inference pipeline on all available data from ADNI and HABS-HD, testing for WM differences across four DTI metrics (FA, MD, RD, AxD), two contrasts (Dementia vs. CN and MCI vs. CN) and two parameterization methods (1D and 2D). Using a large sample size, we aimed to compare 1D and 2D parameterizations and assess whether the additional spatial resolution of 2D parameterization offers substantively different interpretations of WM microstructure, and whether hFDR-RESEL controls false positive detections appropriately relative to the global BH-FDR procedure. Detection rates across metrics, contrasts, parameterizations, and correction methods are summarized in Table 3. Figure 8 shows Cohen’s *d* maps for significant segments or grid cells under hFDR-RESEL (*α*_1_ = 0.05, *α*_2_ = 0.15) for the Dementia vs. CN contrast for the following bundles: the corpus callosum (CC, including the forceps major, forceps minor, precentral and postcentral body, and tapetum), the left cingulum (CG L, including frontal-parietal and parahippocampal portions), the left corticostriatal tracts (CS L, including anterior, precentral, and postcentral portions), the left frontal corticopontine tract (CPT F L), and the left inferior longitudinal fasciculus (ILF L).

**Table 3:** Detection rates (proportion of significant grid cells or segments) across correction methods and profile modes for two contrasts (Dementia vs. CN and MCI vs. CN) and four DTI metrics, aggregated across all bundles. Correction methods applied include global BH-FDR (*α* = 0.05) and hFDR-RESEL (*α*_1_ = 0.05) with *α*_2_ = 0.15 and *α*_2_ = 0.25, shown respectively as hFDR(0.15) and hFDR(0.25).

| Parameterization |  | 1D |  |  | 2D |  |  |
| --- | --- | --- | --- | --- | --- | --- | --- |
| Correction |  | BH-FDR | hFDR(0.15) | hFDR(0.25) | BH-FDR | hFDR(0.15) | hFDR(0.25) |
| Dementia | AxD | 81.3% | 79.4% | 81.6% | 53.2% | 52.6% | 56.8% |
|  | FA | 87.7% | 85.6% | 88.3% | 79.9% | 79.7% | 82.6% |
|  | MD | 93.3% | 92.4% | 93.9% | 58.4% | 58.2% | 62.5% |
|  | RD | 91.5% | 90.6% | 91.7% | 57.6% | 57.4% | 62.1% |
| MCI | AxD | 42.5% | 41.4% | 45.0% | 12.7% | 13.7% | 15.5% |
|  | FA | 56.3% | 53.7% | 60.0% | 38.2% | 42.3% | 49.0% |
|  | MD | 73.9% | 70.7% | 75.4% | 20.2% | 20.7% | 23.5% |
|  | RD | 73.8% | 70.3% | 75.3% | 21.1% | 22.3% | 25.4% |

**Figure 8:**
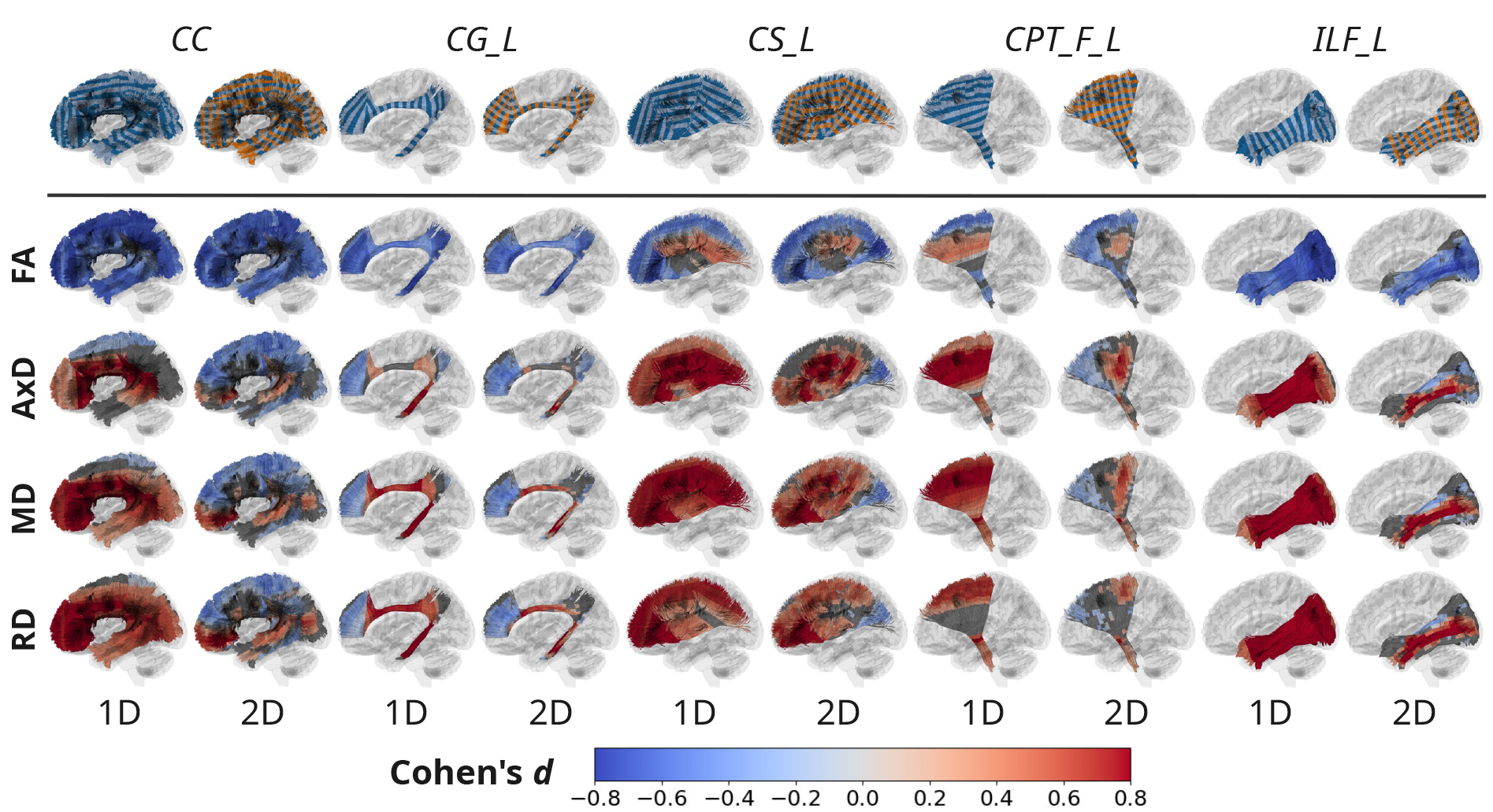
Visualizing Cohen’s *d* for significant results under hFDR-RESEL (*α*_1_ = 0.05, *α*_2_ = 0.15) for the Dementia vs. CN contrast, across 4 DTI metrics (FA, AxD, MD and RD) and 2 modes of parameterization (1D and 2D). The top row plots the 1D segments and 2D grids in alternating colors. For Cohen’s *d* plots, segments or grid cells that were masked or not significant are colored grey. (CC: corpus callosum; CG L: cingulum left; CS L: corticostriatal tract left; CPT F L: corticopontine tract frontal left; ILF L: inferior longitudinal fasciculus left.)

#### Detection rate for 1D versus 2D parameterization

Detection rates are consistently lower in 2D than in 1D across all metrics and contrasts. The magnitude of this reduction differs by metric: FA shows the smallest drop, whereas the MD and RD exhibit far larger reductions. This pattern is likely because 2D parameterization can resolve spatial heterogeneity radially across the bundle, also evident in Figure 8.

#### Detection rate for dementia versus MCI

The reduction in detection rate from 1D to 2D is systematically larger for MCI than Dementia (e.g., 34.9 vs. 53.7 percentage points respectively for MD corrected with BH-FDR). This suggests that MCI-related abnormalities are likely more spatially focal than the diffuse alterations observed in dementia, which is also consistent with the known spread of pathology from MCI to dementia (Feng et al., 2024). Under 2D parameterization, focal effects confined to a sub-region of a segment are preserved instead of averages, but they must survive the larger multiple comparisons burden imposed by the additional number of tests.

#### Detection rate across DTI metrics

With 1D profiling, MD and RD yield the highest detection rate, followed by FA and then AxD for both contrasts, consistently with prior studies (Feng et al., 2024). With 2D profiling, however, FA overtakes MD and RD in detection rate by a larger margin for MCI. The order of the DTI metrics, in terms of sensitivity, is consistent across correction methods.

#### Detection rate across correction methods

Across all conditions, hFDR-RESEL(*α*_2_ = 0.25) achieves the highest detection rate, followed by BH-FDR and hFDR-RESEL(*α*_2_ = 0.15). For 2D parameterization, hFDR-RESEL(*α*_2_ = 0.15) achieves comparable or higher detection than BH-FDR in several conditions, most notably for MCI effect (FA: 42.3% vs. 38.2%; AxD: 13.7% vs. 12.7%).

#### Visualizing dementia effects for FA

In CC, CG L, ILF L, both 1D and 2D reveal a consistent and spatially widespread decrease in FA, confirming that FA reduction in these bundles is largely uniform. In CS L and CPT F L, 2D reveals that the FA increase observed in these bundles is spatially confined to specific sub-regions, instead of spanning the entire cross-section.

#### Visualizing dementia effects for diffusivity metrics

Differences between the two parameterizations are pronounced for the diffusivity metrics. Under 1D profiling, CC shows predominantly increased AxD, MD, and RD in most of the bundle except at the lateral edge segments for AxD; under 2D profiling, however, decreased diffusivity is shown to be more widespread, whereas increased diffusivity is concentrated at the central core. For the forceps minor in particular, we observe decreased diffusivity in the dorsal region and the opposing effect in the ventral region with 2D profiling, but only increased diffusivity with 1D profiling. Such a reversal of the apparent effect direction highlights the hazard of interpreting results from a 1D parameterization when an effect changes radially: averaging over cross-sections yields a mean that reflects the dominant signal and may obscure opposing but weaker signals, an effect compounded by the distance-weighted approach, which weights the bundle core more heavily and further suppresses contributions from the periphery. For the diffusivity metrics in the remaining bundles, the 2D maps provide finer spatial resolution while remaining broadly consistent with the 1D profiles. In the cingulum, we observe decreased diffusivity in the anterior cortical projections, and increased diffusivity in the subgenual and parahippocampal regions. 2D profiling better separates the opposing effects, whereas the striped appearance from 1D likely reflects the segment boundary from along-tract parameterization where the bundle centroid curves sharply. We also observe decreased diffusivity in the posterior portion of CS L, the anterior portion of CPT F L, and the dorsal portion of ILF L, under 2D profiling, patterns that are either absent or ambiguous in the corresponding 1D maps.

Regions with concurrent FA and MD decreases —such as the anterior cortical projection of the cingulum, the edge segments of the corpus callosum —may reflect neuroinflammatory processes resulting in restricted water diffusion and disrupted tissue organization, though this interpretation remains speculative as DTI metrics cannot distinguish among contributing cellular mechanisms.

## 4 Discussion

In this study, we present SPECTRA, a framework for spatially resolved multi-bundle tractometry that integrates 2D bundle parameterization with spatially-aware inference via hierarchical FDR. In synthetic experiments across 144 experimental conditions with varying sample size, effect magnitude and effect geometry, the hFDR-RESEL approach consistently outperformed standard global FDR and hFDR-Bundle, in balancing power and error control. In empirical analyses of MCI and dementia, 2D parameterization revealed spatial patterns of microstructure differences across multiple bundles absent in 1D profiles. In the following sections, we discuss the methodological implications of these findings, and offer practical guidance for designing multi-bundle tractometry studies.

### 4.1 Parameterization and Statistical Inference as a Unified Design

Tractometry occupies a middle ground between voxel-based analysis of diffusion MRI and ROI-based regional summaries: it recovers anatomical interpretability by aggregating measurements along biologically meaningful axes while preserving spatial resolution along the tract. This introduces an important design consideration. The spatial units over which inference is conducted should span approximately equal physical extent, so that effect estimates are on a common scale and findings at different locations carry consistent interpretations within and across bundles. In standard 1D tractometry, each along-tract segment aggregates measures from the full cross section. Because bundle geometry varies along the tract, segments passing through a wide or fanning region summarize substantially larger areas than those in narrower portions. This geometric inconsistency produces heteroscedasticity and complicates interpretation, as statistical findings may reflect variation in bundle geometry rather than the underlying signal. Our 2D parameterization addresses this by constraining each grid cell to a fixed physical extent in both the along-tract and radial dimensions, so that significant cells carry more consistent physical interpretations even across bundles if a common resolution is used.

Building on this parameterization, the hierarchical FDR procedure is designed to account for spatial correlation within each bundle while maintaining error control. We draw on the idea of RESELs from classic RFT as a heuristic for defining block sizing. In RFT, a RESEL is a descriptor of smoothness and an active component of familywise error control. Here, Matérn -derived length scales are used to partition each bundle into spatially contiguous blocks that each capture approximately one unit of spatial information so that Stage 1 aggregation is sensitive. The 2D parameterization makes this particularly well-motivated, as the Matérn kernel can be estimated along anatomically meaningful axes. Importantly, error control in hFDR relies on the validity of the Simes block level statistics and BH procedure under PRDS. This means the procedure remains valid for 1D profiles, although nonstationarity may be more pronounced. The irregularity of 1D segment geometry affects the quality of the smoothness estimate and the precision of block boundaries, but does not threaten the validity of the procedure.

The 2D parameterization exposes spatial structure suppressed by 1D profiling, with direct consequences for interpretation. In 1D, a test at segment *s* is a test of the cross-sectional mean, and a non-significant result could reflect the absence of effect or cancellation of opposing effects. Even a significant result may combine spatially heterogeneous patterns, obscuring their spatial origin. With 2D parameterization, such effects are spatially resolved rather than averaged away. In our empirical results, microstructural differences associated with dementia were often concentrated at the bundle peripheries, patterns that were attenuated or entirely missed by 1D summaries. The corpus callosum illustrates this most clearly, where regions of decreased diffusivity were substantially more spatially extensive in 2D profiles (Figure 8).

Taken together, the parameterization and inference design in SPECTRA are not independent choices. The physical regularity of the 2D grid makes the spatial correlation structure tractable for block-based inference, and the inferential guarantees in turn make the spatially resolved maps interpretable and statistically grounded.

To better understand the effect of RESEL blocking on the residual covariance structure, we examined the eigenvalue spectrum of the noise covariance before and after block-level averaging **see Supplementary section 3**. The eigenvalues of the raw residual covariance decay gradually across hundreds of spatial components, whereas the spectrum decays more steeply with RESEL blocking, indicating that a smaller number of components accounts for the same proportion of total variance and reflecting the suppression of high-frequency spatial noise by the Matérn -derived blocks. This provides an additional way to understand the joint design of parameterization and inference: the 2D parameterization along anatomically meaningful axes helps to estimate the spatial correlation structure more reliably, which in turn determines the block boundaries used in Stage 1 aggregation, and adjusts the covariance representation toward lower effective dimensionality. We note that hFDR-RESEL is designed for detection rather than class separation, and the spectral steepening characterizes the noise covariance specifically rather than the alignment between the noise structure and the disease contrast. Future work could examine parameterization and encoding strategies that more directly bridge these objectives. For instance, adaptive partitioning schemes may better capture spatial correlation within and even across bundles, and approaches that explicitly align the spectral basis with the target effect vector can optimize for discriminability in directions relevant to the disease contrast.

#### A Note on Visualization

In our figures, atlas segments and grid cells are colored uniformly according to their statistical summary for both 1D and 2D parameterizations, to help interpret the spatial locations of results and compare across bundles. This convention should be interpreted carefully, particularly for large segments or cells: color indicates that an effect of the estimated magnitude exists *somewhere* within that spatial unit, not that the effect is uniform throughout. In 2D, where opposing patterns within a cross-section are spatially resolved, this distinction is less consequential. In 1D, a colored segment may reflect a strong focal effect, or a diffuse but modest effect that spans the full segment. When using coarse resolutions, users should treat such maps as evidence of the existence and general location of an effect.

### 4.2 Experimental Design for Multi-Bundle Tractometry Studies

Experimental design for multi-bundle tractometry should balance sensitivity and specificity to have sufficient power to detect effects of interest while controlling false discoveries. The right balance depends on the available sample size, data quality, the hypothesized magnitude and spatial extent of effects, and the spatial resolution of the parameterizations. We discuss each of these factors below and offer practical guidance for study design.

#### Sample Size

Sample size is the most consequential design parameter. In our synthetic experiments, we found that the sample size required to reach 70% sensitivity *N*_70_ varied by nearly five-fold across different effect magnitudes and extents, under the same correction method. Even for large, spatially diffuse effects, more than 200 subjects per group were needed to achieve adequate sensitivity with hFDR-RESEL at *α*_2_ = 0.25. These estimates were obtained when testing with only 20 bundles; studies including more bundles would face a larger multiple testing burden and may require even larger samples.

#### Correction Method

The choice of correction method interacts closely with sample size and the hypothesized effect structure. In our experiments, both hFDR-Bundle and hFDR-RESEL achieve good block-level error control across most settings, except at low sample sizes (*N*_*s*_ = 50). Cell-level FDP inflation was observed in Stage 2, most severely for hFDR-Bundle across all effect geometries, and for hFDR-RESEL under spatially localized effects. In both cases, inflation occurs when the true extent of the effect is smaller than the block size, inferred from noise as a “best guess” for the autocorrelation structure of the signal. When effects are expected to be localized, standard FDR is conservative but maintains good error control; hFDR-RESEL retains a meaningful power advantage even at the more stringent *α*_2_ = 0.15. Among the methods we evaluated, hFDR-RESEL offers the best balance of power and error control, with the parameter *α*_2_ serving as a lever that controls how many false discoveries are tolerated among cells within rejected blocks. This parameter should be set to reflect the user’s prior hypothesis.

#### Grid Resolution

The physical size of grid cells (*δ*_*s*_, *δ*_*r*_) defines the unit of spatial inference and influences both the overall statistical power and error control. In our prior work examining 1D segment resolution (Feng et al., 2025), we found that coarse segments are more data efficient, whereas finer segments are better suited for detecting more localized effects, in line with the matched filter theorem. As such, we expect the same qualitative tradeoff to hold with 2D parameterizations. Cells that are too small risk sparse streamline coverage, unstable LMM fits, and unreliable Matérn length scale estimates, and cells that are too large may obscure local effects. For 2D parameterization specifically, *δ*_*s*_ should not be set too large, as the radial dimension is derived via PCA within each along-tract segment. This assumes the local streamline geometry is approximately linear within a segment, an assumption that becomes less tenable as segments span regions of greater curvature (Yue et al., 2016). In SPECTRA, users may supply *δ*_*s*_ and *δ*_*r*_ directly when generating profiles, and should select values appropriate to their data and scientific question. An important consideration in multi-bundle analysis is that grid resolution should be kept consistent across bundles, so that the number of statistical tests is approximately proportional to the physical volume of tissue examined rather than reflecting arbitrary differences in parameterization choices. Future work will characterize the sensitivity and error control implications of grid resolution and how they interact with sample size, dataset quality and variability. We provide the synthetic effect injection module in SPECTRA for users who wish to conduct such analyses on their own data.

#### Parameterization Mode and Bundle Geometry

The choice between 1D and 2D parameterization should be informed by bundle geometry. For bundles with small, geometrically consistent cross sections, such as the fornix and parahippocampal cingulum, 1D tractometry is likely sufficient and provides greater statistical power by reducing the number of statistical tests. For bundles with large cross-sections, 2D parametrization is preferable as it resolves spatial heterogeneity that 1D averaging would suppress. Importantly, mixing 1D and 2D parameterizations in the same analysis creates imbalance in the number of tests contributed per bundle, although hFDR still remains valid. Consistent grid resolution makes multi-bundle inference interpretable, and is best achieved by committing to a single mode.

Atlas bundle definitions are typically derived from anatomical priors and population averages, and primarily serve as references for segmentation. For tractometry, additional refinement may be beneficial. For 1D analysis, a bundle with large cross-sections or bifurcating geometry may be subdivided to improve along-tract consistency. For 2D analysis, bundles with sharp curvature, e.g., the cingulum and SLF-I, can produce small grid cells in high curvature regions; SPECTRA addresses this through subject-level masking to avoid making inferences at these locations. Even so, such bundles warrant careful inspection. Bundles with highly variable thickness or complex branching geometry, e.g., SLF-II, may require more advanced geometric modeling beyond what 2D parameterizations can accommodate (Shailja et al., 2023). Such advanced parameterization would also require strict alignment between subject and atlas, and streamline-based registration such as BundleWarp (B. Q. Chandio et al., 2023) may be used to further improve correspondence.

#### Data Quality

Data quality affects many aspects of dMRI processing, including tractography, diffusion model fitting, and by extension, tractometry (Schilling et al., 2021). In regions where atlas bundles have full coverage but individual subjects fail to reconstruct, a robust parameterization method should accommodate incomplete or missing data. Our 2D parameterization follows BUAN (B. Q. Chandio et al., 2020), which defines the segments on the atlas, so that missing subject data at a given spatial location does not propagate to neighboring segments. SPECTRA applies subject-level masking based on streamline point count, followed by a group-level coverage threshold to ensure that statistical tests are conducted only where sufficient data exist. Users may adjust this threshold to suit their data. Even so, quality control of bundle reconstructions is important to ensure valid statistical testing. Depending on the data quality, regional variation in tractography means that the effective sample sizes at individual grid cells may be lower, particularly at bundle extremities with poor coverage.

## 5 Conclusion

We introduce SPECTRA, a framework for spatially resolved mapping of white matter microstructure that integrates 2D bundle parameterization with spatially adaptive hierarchical FDR inference. The proposed hFDR procedure improves statistical power and data efficiency by aggregating evidence at coarser block levels prior to finer cell-level testing, with data-driven RESEL blocks defined via a Matérn kernel. In extensive simulations, the hFDR-RESEL approach achieved substantial gains in power while maintaining comparable error control, particularly for spatially diffuse effects, relative to standard FDR approaches. We further demonstrate SPECTRA in an empirical study of cognitive decline in a large multi-site cohort, where 2D parameterization reveals radially structured microstructural differences across multiple bundles, including opposing effects within the same cross-section that are entirely obscured by conventional along-tract profiling. Together, these results suggest that spatially resolved parameterization and adaptive correction procedures are complementary: the 2D grid exposes structure that motivates more targeted inference, and hFDR exploits such spatial structure to improve sensitivity. Overall, SPECTRA is designed to enhance both anatomical precision and statistical sensitivity in tractometry analyses. As neuroimaging datasets continue to grow in scale and complexity, such approaches will be increasingly important to achieve fine-grained and interpretable insights into WM organization.

## Data and Code Availability

The SPECTRA Python package is openly available at https://github.com/wendyfyx/SPECTRA, and includes command-linear interfaces for tractometry profiling, statistical inference, and visualization. The white matter atlas used for bundle tractography is available at https://doi.org/10.5281/zenodo.19656193. Data derivatives, including tractometry profiles, are available to researchers with appropriate data use agreements for the Alzheimer’s Disease Neuroimaging Initiative (ADNI) and the Health & Aging Brain Study - Health Disparities (HABS-HD) datasets upon request; access may be requested through the Image and Data Archive (IDA; https://ida.loni.usc.edu).

## Author Contributions

Y.F.: writing - original draft, writing - review & editing, conceptualization, methodology, software, formal analysis, investigation, resources, data curation, visualization; J.E.V.: data curation, resources, investigation, writing - review & editing; I.B.G.: data curation, resources, writing - review & editing; J.D.A.: data curation, resources; S.I.T.: data curation, investigation, project administration; K.L.: data curation, investigation; S.S.: data curation, investigation; H.Y.: data curation; Y.S.: data curation; S.C.: data curation; T.M.N.: data curation, resources, investigation, writing - review & editing, supervision; N.J.: writing - review & editing, supervision, funding acquisition; B.Q.C.: conceptualization, data curation, resources, writing - review & editing, supervision; P.M.T.: conceptualization, writing - review & editing, supervision, funding acquisition.

## Ethics

Informed consent was obtained for all participants in the ADNI and HABS-HD datasets.

## Acknowledgements

This research was supported by the National Institutes of Health (NIH) grants U01 AG068057, RF1 NS136995, RF1 AG057892, S10 OD032285 and the Alzheimer’s Association grant AARG-23-1150420.

Data collection and sharing for this project was funded by the Alzheimer’s Disease Neuroimaging Initiative (ADNI) (National Institutes of Health Grant U01 AG024904) and DOD ADNI (Department of Defense award number W81XWH-12-2-0012). ADNI is funded by the National Institute on Aging, the National Institute of Biomedical Imaging and Bioengineering, and through generous contributions from the following: AbbVie, Alzheimer’s Association; Alzheimer’s Drug Discovery Foundation; Araclon Biotech; BioClinica, Inc.; Biogen; Bristol-Myers Squibb Company; CereSpir, Inc.; Cogstate; Eisai Inc.; Elan Pharmaceuticals, Inc.; Eli Lilly and Company; EuroImmun; F. Hoffmann-La Roche Ltd and its affiliated company Genentech, Inc.; Fujirebio; GE Healthcare; IXICO Ltd.; Janssen Alzheimer Immunotherapy Research & Development, LLC.; Johnson & Johnson Pharmaceutical Research & Development LLC.; Lumosity; Lundbeck; Merck & Co., Inc.; Meso Scale Diagnostics, LLC.; NeuroRx Research; Neurotrack Technologies; Novartis Pharmaceuticals Corporation; Pfizer Inc.; Piramal Imaging; Servier; Takeda Pharmaceutical Company; and Transition Therapeutics. The Canadian Institutes of Health Research is providing funds to support ADNI clinical sites in Canada. Private sector contributions are facilitated by the Foundation for the National Institutes of Health (www.fnih.org). The grantee organization is the Northern California Institute for Research and Education, and the study is coordinated by the Alzheimer’s Therapeutic Research Institute at the University of Southern California. ADNI data are disseminated by the Laboratory for Neuro Imaging at the University of Southern California.

Generative AI tools were used during the preparation of this work. For the accompanying SPECTRA software, AI tools supported code refactoring, debugging, and documentation. For the manuscript, AI tools were used for grammatical checks and text restructuring for clarity. All ideas, study design, methodological contributions, analyses, interpretations, and prompts directing the tools are entirely original to the authors. All AI-assisted outputs have been carefully reviewed, tested where applicable, and edited by the authors, who take full responsibility and remain fully accountable for all content in this submission.

