## Supplementary Materials for "SPECTRA: Spatial Inference for Tractometry Toward Precision Mapping of White Matter Microstructure"

May 5, 2026

#### Contents


**1 List of Bundles** **2**

**2 Synthetic Effects Evaluation for 1D Tractometry** **4**

**3 Residual Covariance Spectrum Analysis** **7**

### 1 List of Bundles

The following is a list of 63 bundles adapted from the HCP1065 atlas used in this study. All bundles were used in the empirical experiments; the subset of 20 bundles used for synthetic experiments are marked with a checkmark. Radius and max angle indicate values of bundle-specific tracking parameters. Except for the commissural bundles, all other bundles are bilateral, where each of the left and right bundles were extracted and analyzed separately.

| Abbreviation | Full Name | Synthetic | Radius | Max Angle |
|---|---|---|---|---|
| **Association** | | | | |
| AF | Arcuate Fasciculus | ✓ | 6 | 20 |
| CG_FP | Cingulum Frontal-Parietal | ✓ | 6 | 20 |
| CG_PH | Cingulum Parahippocampal | | 6 | 20 |
| CG_PO | Cingulum Parolfactory | | 6 | 25 |
| CG_SLF1 | Cingulum Superior Longitudinal Fasciculus 1 | | 6 | 20 |
| EMC | Extreme Capsule | | 7 | 20 |
| FAT | Frontal Aslant Tract | | 6 | 20 |
| HA | Hippocampus Alveus | | 6 | 20 |
| IFOF | Inferior Fronto-Occipital Fasciculus | ✓ | 7 | 20 |
| ILF | Inferior Longitudinal Fasciculus | | 6 | 20 |
| MdLF | Middle Longitudinal Fasciculus | | 7 | 20 |
| PAT | Parietal Aslant Tract | | 6 | 20 |
| SLF2 | Superior Longitudinal Fasciculus 2 | | 6 | 20 |
| SLF3 | Superior Longitudinal Fasciculus 3 | | 6 | 20 |
| UF | Uncinate Fasciculus | ✓ | 6 | 25 |
| VOF | Vertical Occipital Fasciculus | | 6 | 20 |
| **Commissure** | | | | |
| CC_Body_Post | Corpus Callosum Body Postcentral | ✓ | 8 | 20 |
| CC_Body_Pre | Corpus Callosum Body Precentral | ✓ | 8 | 20 |
| CC_ForcepsMajor | Corpus Callosum Forceps Major | ✓ | 8 | 20 |
| CC_ForcepsMinor | Corpus Callosum Forceps Minor | ✓ | 8 | 20 |
| CC_Tapetum | Corpus Callosum Tapetum | | 8 | 20 |
| **Projection (Basal Ganglia)** | | | | |
| AR | Acoustic Radiation | | 6 | 20 |
| CS_A | Corticostriatal Tract Anterior | | 6 | 20 |

| Abbreviation | Full Name | Synthetic | Radius | Max Angle |
|---|---|---|---|---|
| CS_Post | Corticostriatal Tract Postcentral | | 6 | 20 |
| CS_Pre | Corticostriatal Tract Precentral | | 6 | 20 |
| FX | Fornix | | 6 | 30 |
| OR | Optic Radiation | | 6 | 30 |
| TR_A | Thalamic Radiation Anterior | ✓ | 6 | 20 |
| TR_Post | Thalamic Radiation Postcentral | ✓ | 6 | 20 |
| TR_Pre | Thalamic Radiation Precentral | ✓ | 6 | 20 |
| **Projection (Brainstem)** | | | | |
| CPT_F | Corticopontine Tract Frontal | | 6 | 25 |
| CPT_P | Corticopontine Tract Parietal | | 6 | 25 |
| CST | Corticospinal Tract | ✓ | 6 | 20 |
| ML | Medial Lemniscus | | 6 | 20 |

### 2 Synthetic Effects Evaluation for 1D Tractometry

The simulation experiments were repeated on 1D tractometry profiles with the same 144 experimental conditions as those used for the 2D profiles. The block- and cell-level sensitivity and false discovery proportion (FDP) are shown in Figure 1 and Figure 2.

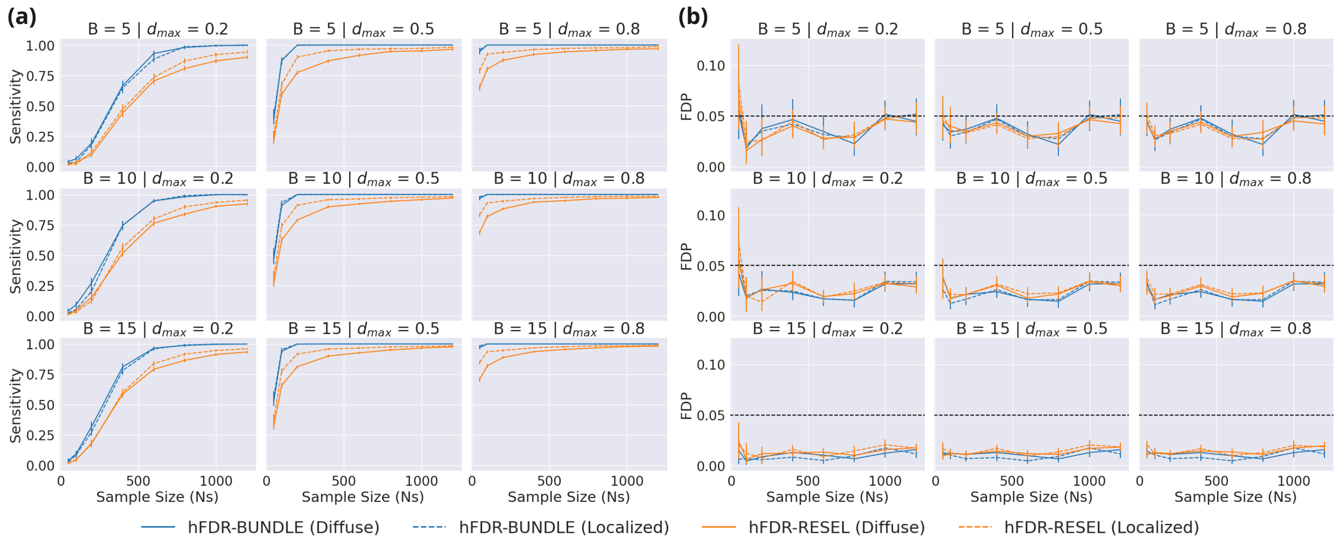


Figure 1: Block-level evaluation for simulation experiments for two variants of hFDR for 1D tractometry profiles. Sensitivity (a) and FDP (b) for each experimental condition is shown with increasing sample sizes $N_s$ across different peak effect sizes ($d_{\max}$) and numbers of affected bundles ($B$). The nominal $\alpha = 0.05$ level is shown in panel (b) as a black dashed line.

The overall qualitative ordering of methods is preserved compared to the 2D results. At the block level, hFDR-Bundle achieves higher sensitivity than hFDR-RESEL, and FDP for both variants remains below the nominal $\alpha = 0.05$ across all conditions. At the cell level, the ranking of correction methods in both sensitivity and FDP mirrors the 2D pattern, with hFDR-Bundle most sensitive but least conservative, hFDR-RESEL striking a favorable balance, and the global BH procedures the most conservative.

Two patterns differ from the 2D results, both likely attributable to the smaller grids produced by 1D parameterization. First, the the block-level sensitivity gap between diffuse and localized effects is markedly smaller in 1D for both hFDR variants (Figure 1). Second, the cell-level sensitivity advantage of hFDR-RESEL over the global BH procedures shrinks with increasing $d_{\max}$ and number of affected bundles $B$ (Figure 2). The same trend is present in 2D, but in 1D the advantage collapses almost entirely at large $d_{\max}$ across both effect geometries. With far fewer along-tract segments per bundle, the two effect geometries are less distinguishable compared to their 2D counterparts. Under 1D, the null fraction is lower to begin with, and the global BH procedure can adequately capture the effect when the signal is

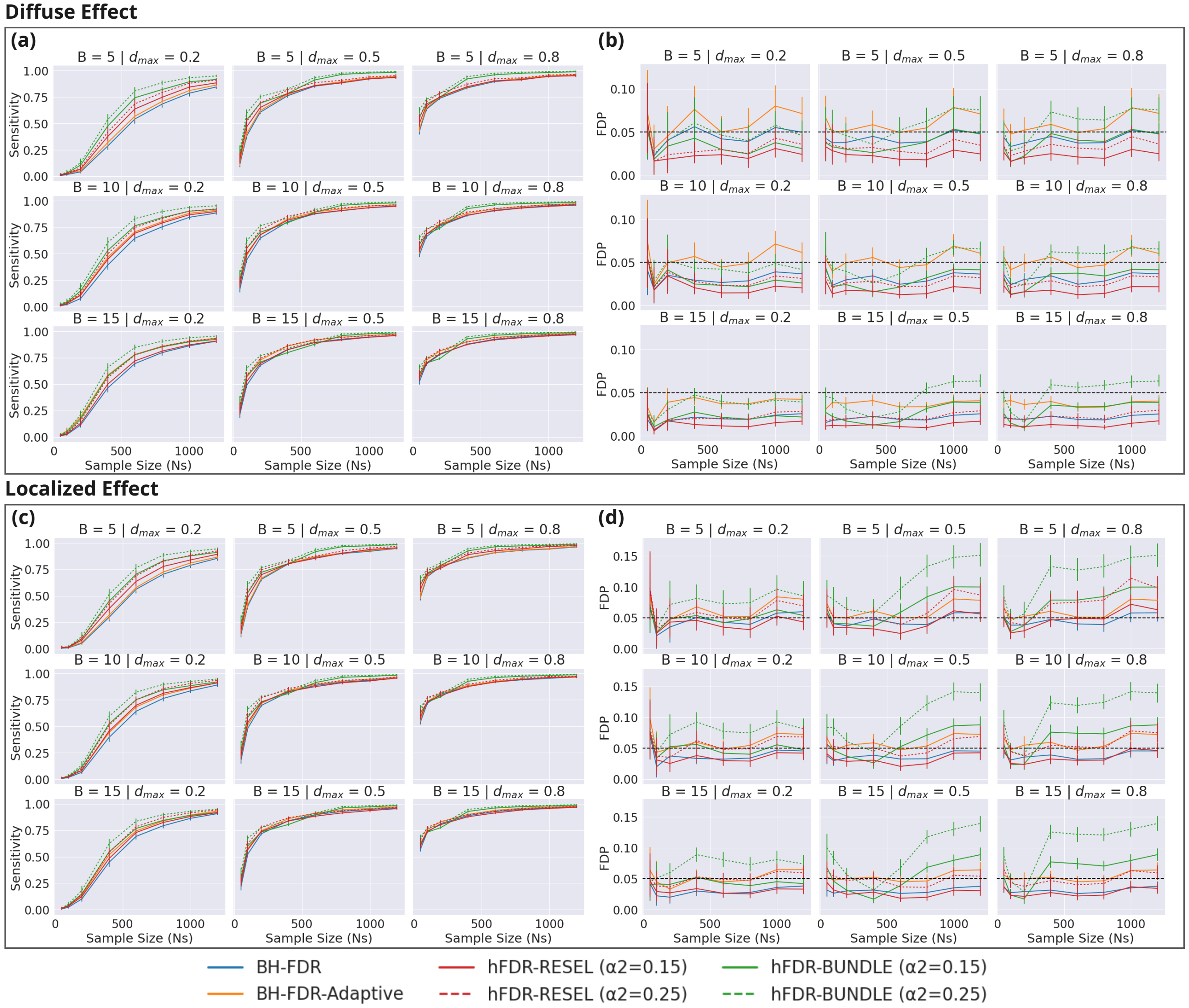


Figure 2: Cell-level evaluation for simulation experiments across 4 correction procedures, with different $\alpha_2$ shown as different line styles for the hFDR variants for 1D tractometry profiles. Sensitivity (a, c) and FDP (b, d) for each experimental condition is plotted with increasing sample sizes $N_s$ across different peak effect sizes ($d_{\max}$) and numbers of affected bundles ($B$), separately for diffuse and localized effects. The nominal $\alpha = 0.05$ level is shown in panel (b, d) as a horizontal black dashed line.

strong enough.

Overall, the global BH procedures remain effective for 1D profiles across many experimental conditions, with hFDR-RESEL offering a meaningful sensitivity advantange primarily at low effect sizes and small $B$, while maintaining valid error control.

### 3 Residual Covariance Spectrum Analysis

To characterize the effect of RESEL block averaging on the spatial noise structure, we computed the eigenvalue spectrum of the residual covariance matrix before and after blocking for a representative bundle. Reduced-model residuals, obtained by fitting the LMM with covariates only, excluding the target variable, were extracted at all valid grid cells, yielding a matrix of shape $(N_{\text{subj}} \times N_{\text{cells}})$. The sample covariance was decomposed via singular value decomposition (SVD), giving eigenvalues $\lambda_i = \sigma_i^2/(N_{\text{subj}} - 1)$ in descending order; near-zero eigenvalues ($\lambda_i < 10^{-12}$) were discarded. Eigenvalues were normalized to sum to one. A power-law decay $\log \lambda_i = -\beta \log i + c$ was fit by ordinary least squares on the top half of eigenvalues to avoid the noise floor at high ranks, yielding the spectral slope $\beta$.

For the RESEL-encoded representation, the same reduced-model residuals were averaged within each RESEL block using the Matérn length scales estimated from the reduced-model residuals. This produced a matrix of shape $(N_{\text{subj}} \times N_{\text{blocks}})$, to which the same SVD and power-law fitting procedure was applied. The steeper slope after RESEL blocking ($\beta = 1.48$ vs $\beta = 1.16$ for CST_L, see Figure 3 *left*; $\beta = 1.31$ vs $\beta = 1.12$ for CC_ForcepsMajor, see Figure 4 *left*) indicates that residual variance is more strongly concentrated in the leading components.

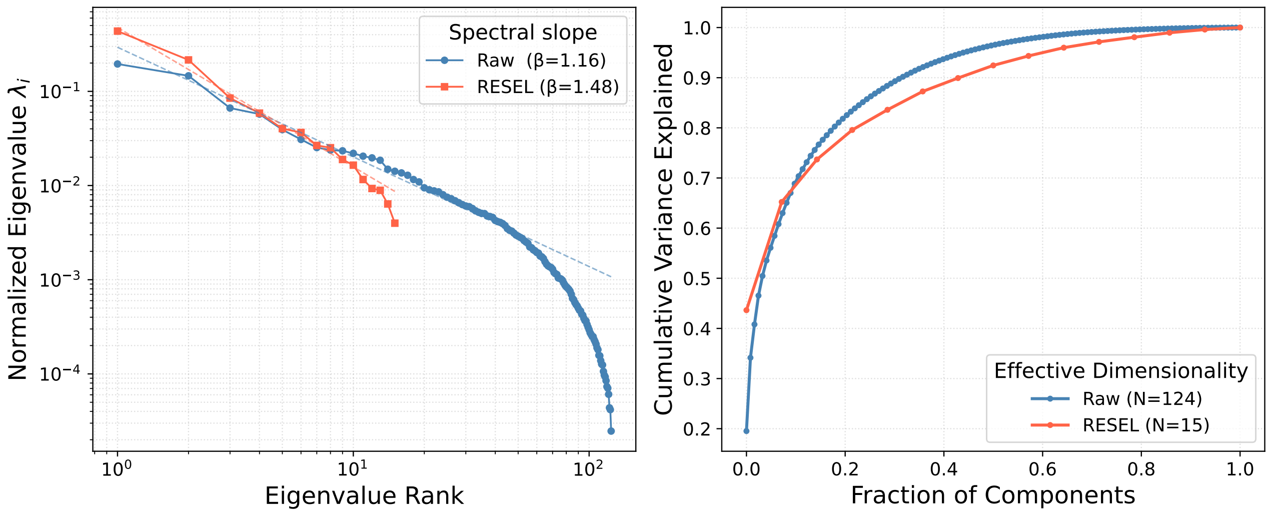


Figure 3: Residual covariance spectrum before and after RESEL block averaging for CST_L where the model was fitted on 2D profiles of FA on the full available sample. **Left:** Log-log plot of normalized eigenvalues $\lambda_i$ of the residual covariance matrix, ranked in descending order, for raw grid cells $(N = 250)$ and RESEL block-averaged residuals $(N = 23)$. Dashed lines show power-law fits to the top half of eigenvalues. **Right:** Cumulative variance explained as a function of the fraction of components retained, with the x-axis expressed as fractional rank to allow direct comparison between the raw and RESEL representations. This reflects the reduction in effective dimensionality from 124 valid grid cells to 15 RESEL blocks.

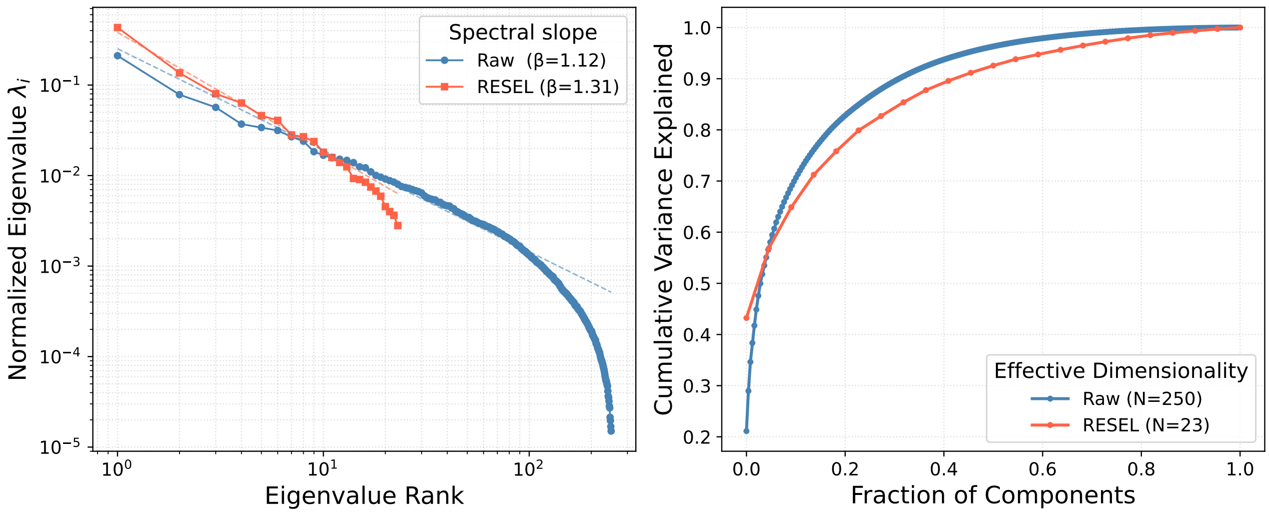


Figure 4: Residual covariance spectrum before and after RESEL block averaging for CC_ForcepsMajor where the model was fitted on 2D profiles of FA on the full available sample.

We note that averaging residuals within RESEL blocks here serves as an approximate encoding strategy to characterize the covariance structure of the blocked representation in hFDR Stage 1, but does not correspond directly to the hFDR inference procedure. In hFDR, Stage 1 aggregation is performed via Simes' test applied to cell-level $p$-values within each block, and detection is formalized through conditional $p$-values at Stage 2. This analysis is meant to characterize how RESEL block boundaries shape the spatial noise covariance, rather than a direct account of the mechanism underlying hFDR's power gains.
